# Heteromeric LGP2-MDA5 receptor complexes drive antiviral signaling

**DOI:** 10.64898/2026.08.20.745998

**Authors:** Gina Buchel, Ling Xu, Yu-Chieh Sung, Anna Marie Pyle

## Abstract

The innate immune receptor MDA5 serves as a first line of defense against RNA viruses by recognizing viral double-stranded RNA (dsRNA) and initiating antiviral signaling. MDA5 is upregulated by another RIG-I-like receptor (RLR) family member known as LGP2, which has been shown to enhance MDA5 activity by recruiting MDA5 to dsRNA. However, it has remained unclear whether LGP2 functions solely as a transient recruitment factor or actively participates in signaling-competent MDA5 filaments. Here, we demonstrate that LGP2 does more than recruiting MDA5 to dsRNA, and that LGP2 becomes an integral component of MDA5 filaments. Mechanistically, LGP2 stabilizes MDA5 filaments and reduces the number of MDA5 molecules required to nucleate stable signaling-competent filaments. This interplay between the two RLR proteins is mediated by specific dsRNA-dependent interactions between the C-terminal tail of LGP2 and MDA5, resulting in a network of contacts that is essential for LGP2-mediated enhancement of MDA5 signaling. Together, these findings provide new mechanistic insight into how LGP2 regulates the assembly and persistence of signaling-competent MDA5 filaments, establishing filament architecture and stability as key determinants of MDA5 signaling. Collectively, this work provides a mechanistic framework for understanding LGP2-mediated regulation of MDA5 signaling and advances our knowledge of the molecular mechanisms governing antiviral immunity.

## Introduction

Upon viral infection and cytosolic introduction of pathogenic RNA, a collection of cytoplasmic pattern-recognition receptors (PRRs) within the host cell activate signaling cascades that produce an effective antiviral and inflammatory response (1). Some of the most prevalent PRRs are the RIG-I-like receptors (RLRs), which include three family members: RIG-I (retinoic acid inducible gene I)(2), MDA5 (melanoma differentiation-associated gene 5)(3), and LGP2 (laboratory of genetics and physiology 2)(4). The RLRs share a similar protein core consisting of a central RNA helicase-like motor domain, specialized insertion domains, and a C-terminal regulatory domain (CTD) (5–7). RIG-I and MDA5 also contain a set of caspase recruitment domains (CARDs) that become exposed and capable of signaling upon binding double-stranded RNA (dsRNA). This results in the formation of interactions between the CARDs and an adaptor protein, the mitochondrial antiviral signaling protein (MAVS)(8), resulting in activation of a downstream signaling cascade that induces expression of type-1 and type-3 interferons (IFNs) and proinflammatory cytokines (9, 10). Of the three proteins in the RLR family, LGP2 has long been mysterious because it lacks CARDs and cannot interact directly with MAVS (6), yet it has been shown to regulate RIG-I and MDA5 signaling (11–16).

The RLRs have different dsRNA substrate specificities, enabling them to provide vertebrates with overlapping strategies for recognizing diverse RNA viruses (7, 9). For instance, RIG-I recognizes short blunt-end RNA duplexes with 5’-tri or 5’-di phosphate termini (17–19). MDA5 signaling, on the other hand, is limited to long dsRNAs around 2 kb and longer (19, 20) and MDA5 oligomerizes on dsRNA to form signaling-competent filaments (21–24). In contrast, LGP2 does not require distinct structural features or specific dsRNA lengths, although it binds dsRNA termini with high affinity (25–27), making LGP2 a particularly versatile dsRNA recognition element (16, 25–27). Growing evidence suggests that LGP2 can bind short dsRNA and recruit MDA5 to dsRNAs that would otherwise be suboptimal for MDA5 signaling (27, 28), resulting in a greater number of shorter filaments that can participate in signaling (12, 29, 30).

Although the precise mechanistic role of LGP2 remained elusive for many years, significant advances have been made within the last decade to elucidate how LGP2 promotes MDA5-mediated antiviral signaling. Numerous studies have demonstrated that LGP2 enhances MDA5 dsRNA binding and filament assembly and, in turn, increases signal transduction and overall immune response *in cellulo* (12, 13, 16, 29–32). More specifically, LGP2 was shown to recruit MDA5 to dsRNA and promote the formation of shorter filaments capable of signaling (12, 29, 33). Recent studies showed that ATP-dependent translocation of LGP2 along dsRNA regulates MDA5 filament assembly and antiviral signaling (29, 30, 33). Importantly, recent cryogenic electron microscopy (cryo-EM) studies of the LGP2-MDA5-dsRNA signaling unit have contributed valuable insight into the molecular interactions between LGP2 and MDA5 on dsRNA (31). Despite these advances, important questions remain regarding the stoichiometry, composition, and stability of MDA5 filaments in the presence of LGP2.

Here, we use a combination of cellular, biochemical, and biophysical approaches to further examine the assembly, architecture, stability, and function of MDA5 filaments in the absence and presence of LGP2. Our results provide evidence that LGP2 and MDA5 can coexist on dsRNA, forming a stable complex in which LGP2 molecules remain bound to dsRNA during and after MDA5 filament assembly, indicating that LGP2 functions as more than a transient dsRNA recruitment factor. We further find that LGP2 stabilizes MDA5-dsRNA complexes and reduces the number of MDA5 molecules required for maintaining stable filaments. In addition, our data support that the C-terminal tail (CTT) of LGP2 promotes MDA5 recruitment to dsRNA, facilitates heteromeric complex assembly through specific interaction networks, and enhances MDA5 oligomerization, therefore establishing the CTT as a critical LGP2-MDA5 interaction interface. Together, these findings extend current models of LGP2-dependent MDA5 regulation and provide new mechanistic insight into how LGP2 enhances MDA5 signaling.

## Results

### MDA5 signaling is selectively enhanced by LGP2

To better understand the mechanism by which LGP2 enhances MDA5 signaling *in cellulo*, we used a well-established cell-based reporter system using HEK293T cells, which exhibit low endogenous RLR signaling activity (12). Cells were transfected with plasmids encoding MDA5, LGP2, and/or a stimulatory dsRNA, enabling us to selectively monitor the influence of the RLR or dsRNA molecule of interest. Transfection of the reporter system with MDA5 and a dsRNA of 150 bp (dsRNA150) produced robust interferon signaling (green bar, **Fig. 1A** and **Fig. S1A**), consistent with previous reports demonstrating that MDA5 is activated by sufficiently long synthetic dsRNA molecules (19, 23, 31). When low amounts of LGP2 were also co-expressed, we observed a striking increase in the level of interferon induction (orange bars, **Fig. 1A**). At much higher concentrations of LGP2 relative to MDA5, we observed an eventual reduction in signaling (orange bars, **Fig. 1A**), suggesting that LGP2 begins to compete with MDA5 along the dsRNA lattice. This experiment suggests that enhancement of MDA5 signaling by LGP2 is concentration-dependent, corroborating observations from previous studies (11–13, 34, 35).

**Figure 1.**
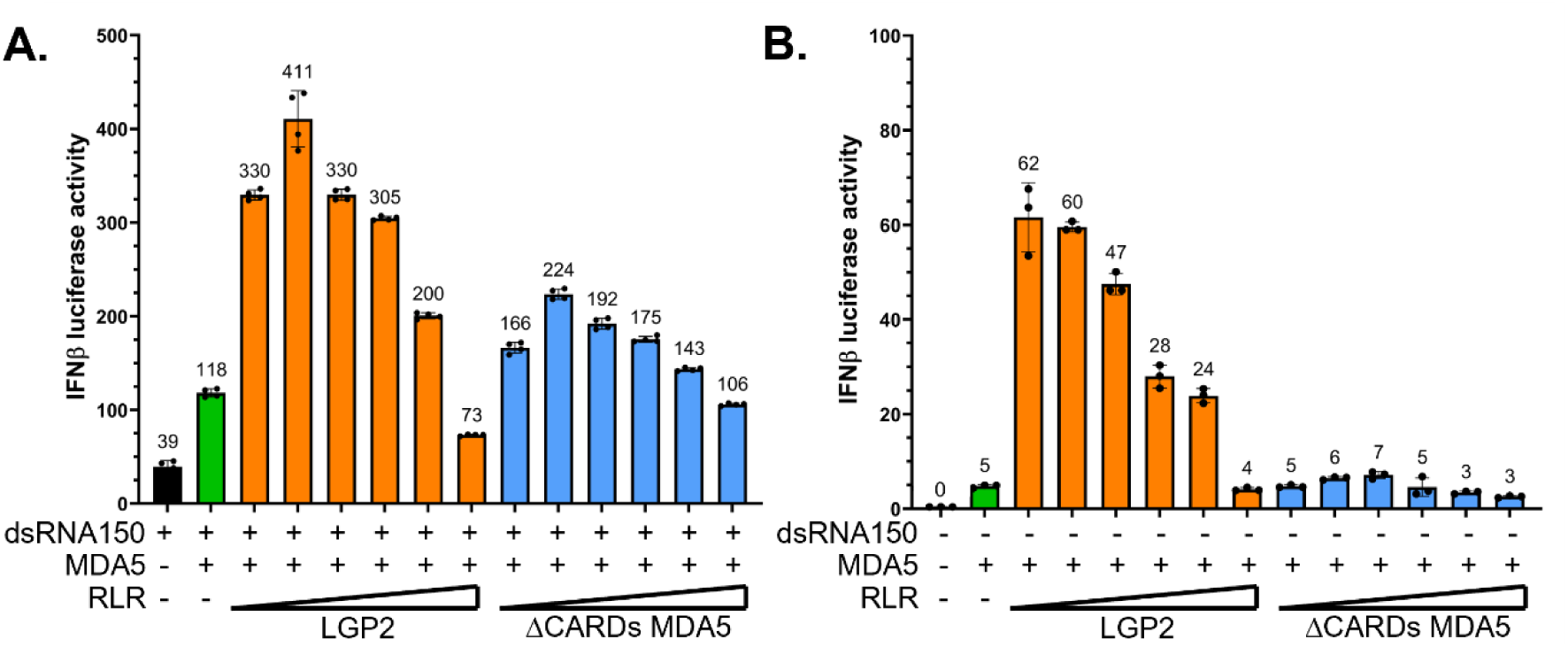
MDA5-mediated signaling is selectively enhanced by LGP2. **A**. IFN induction by co-transfection of MDA5 (1 ng) and LGP2 (orange) or ΔCARDs MDA5 (blue) plasmids (5, 10, 20, 50, 100, 200 ng) stimulated with 1 ug dsRNA150 in HEK293T cells. Data are represented as mean±SD (n=4 biological replicates). **B**. IFN induction by co-transfection of MDA5 (1 ng) and LGP2 (orange) or ΔCARDs MDA5 (blue) plasmids (5, 10, 20, 50, 100, 200 ng) in HEK293T cells. Data are represented as mean±SD (n=3 biological replicates).

Given that LGP2 and MDA5 are structurally similar (6, 27, 36–38), we next investigated whether LGP2 enhances MDA5 signaling simply by adding another subunit of protein to the nascent filament or by contributing a unique function to the MDA5-dsRNA signaling complex. Because LGP2 lacks CARDs and cannot signal independently (6), we asked whether its enhancement effect could be simulated by replacement with a signaling-incompetent MDA5 subunit. To test this “simple subunit” model, we deleted the CARDs from MDA5 (ΔCARDs MDA5) and co-transfected this deletion mutant with full-length MDA5 and dsRNA150, as we had done with LGP2. We found that introduction of excess ΔCARDs MDA5 to full-length MDA5 increases interferon signaling at all concentrations (blue bars, **Fig. 1A**), relative to MDA5 alone (green bar, **Fig. 1A**), but ΔCARDs MDA5 has a weaker stimulatory effect than LGP2. These results indicate that LGP2 contributes a unique function to the MDA5-dsRNA complex that a CARDs-deficient MDA5 cannot recapitulate. While establishing that LGP2 brings something special to the MDA5 filament, our studies with the ΔCARDs MDA5 mutant also have surprising implications for the MDA5 signaling mechanism. These results show that functional MDA5 signaling complexes need not be coated with CARDs, instead requiring only a few CARDs to productively engage MAVS and stimulate signaling on long dsRNAs, challenging conventional tetrad models of CARD-CARD assemblies in which each MDA5 contributes a CARD to the complex (23, 39, 40). In this regard, MDA5 signaling appears more similar than expected to RIG-I signaling, in which the tandem CARDs from a single RIG-I molecule can nucleate MAVS filament formation to initiate the signaling cascade (41).

The definitive advantage of LGP2 over ΔCARDs MDA5 is observed across multiple dsRNA substrates, including low molecular weight (LMW) poly(I:C) and longer synthetic mixed-sequence dsRNA molecules, such as a dsRNA of 300 bp (dsRNA300) (**Fig. S1B,C**). Perhaps most strikingly, when cells are not transfected with dsRNA and only endogenous cellular dsRNAs are present, there was minimal immune activation by MDA5 alone (green bar, **Fig. 1B**). But this abruptly changed when LGP2 was co-expressed, and robust signaling was observed (orange bars, **Fig. 1B**), presumably stimulated by endogenous dsRNA molecules. In contrast, we did not observe any enhancement of MDA5 signaling upon addition of ΔCARDs MDA5 in the absence of exogenous RNA (blue bars, **Fig. 1B**). This result indicates that only LGP2 allows MDA5 to signal on short, endogenous dsRNAs from the host, suggesting the LGP2-MDA5 interface contributes distinct functions relative to the MDA5-MDA5 interface. Taken together, these data demonstrate that LGP2 possesses features that enable the protein to specifically enhance MDA5 signaling on a diversity of dsRNAs, emphasizing the importance of examining attributes of the heteromeric LGP2-MDA5 complex.

### MDA5 and LGP2 form a stable heteromeric complex on synthetic dsRNA

To evaluate the properties of the heteromeric LGP2-MDA5-dsRNA150 complex *in vitro*, we conducted electrophoretic mobility shift assays (EMSAs) in which either dsRNA or proteins (MDA5 or LGP2) were individually labeled with ^32^P using a kinase. We first sought to determine if the heteromeric LGP2-MDA5 complex can be assembled *in vitro* on radiolabeled dsRNA150 (dsRNA150-^32^P) and to evaluate whether both proteins bind dsRNA simultaneously. Since LGP2 has a faster on-rate compared to MDA5 (12, 25), we first loaded LGP2 onto dsRNA150. We observed a shift in band migration when LGP2 was incubated with dsRNA150-^32^P, representing the formation of a discrete LGP2-dsRNA150 complex (**Fig. 2A**, Lanes 1,2). This result indicates that LGP2 forms a stable complex with dsRNA150, predominantly containing one or two bound LGP2 molecules (**Fig. S2A**). Similarly, MDA5 formed stable complexes with dsRNA150 (**Fig. S2B**). However, the number of MDA5 molecules bound could not be accurately determined, as EMSA lacks the resolution to distinguish filament stoichiometries. When MDA5 was added to the pre-formed LGP2-dsRNA150 complex, a “super-shifted” band appeared (**Fig. 2A**, Lanes 3-6), indicating a higher-order LGP2-MDA5-dsRNA150 complex. We then sought to determine whether LGP2 was still part of this complex by radiolabeling LGP2 with ^32^P (LGP2-^32^P) and leaving dsRNA150 unlabeled (**Fig. 2A**, Lanes 7-12). In this case, we observed that LGP2 remains bound to dsRNA150 even after MDA5 joins the complex, as demonstrated by the accumulation of a super-shifted band (**Fig. 2A**, Lanes 9-12). Finally, to directly monitor the presence of MDA5, we radiolabeled MDA5 (MDA5-^32^P) while leaving dsRNA150 and LGP2 unlabeled (**Fig. 2B**, Lanes 7-12). In this case, we also observed accumulation of a signal coming from MDA5-^32^P bound to LGP2-dsRNA150 with increasing amounts of the radiolabeled protein (**Fig. 2B**, Lanes 9-12), indicating formation of a dsRNA complex that contains both MDA5 and LGP2.

**Figure 2.**
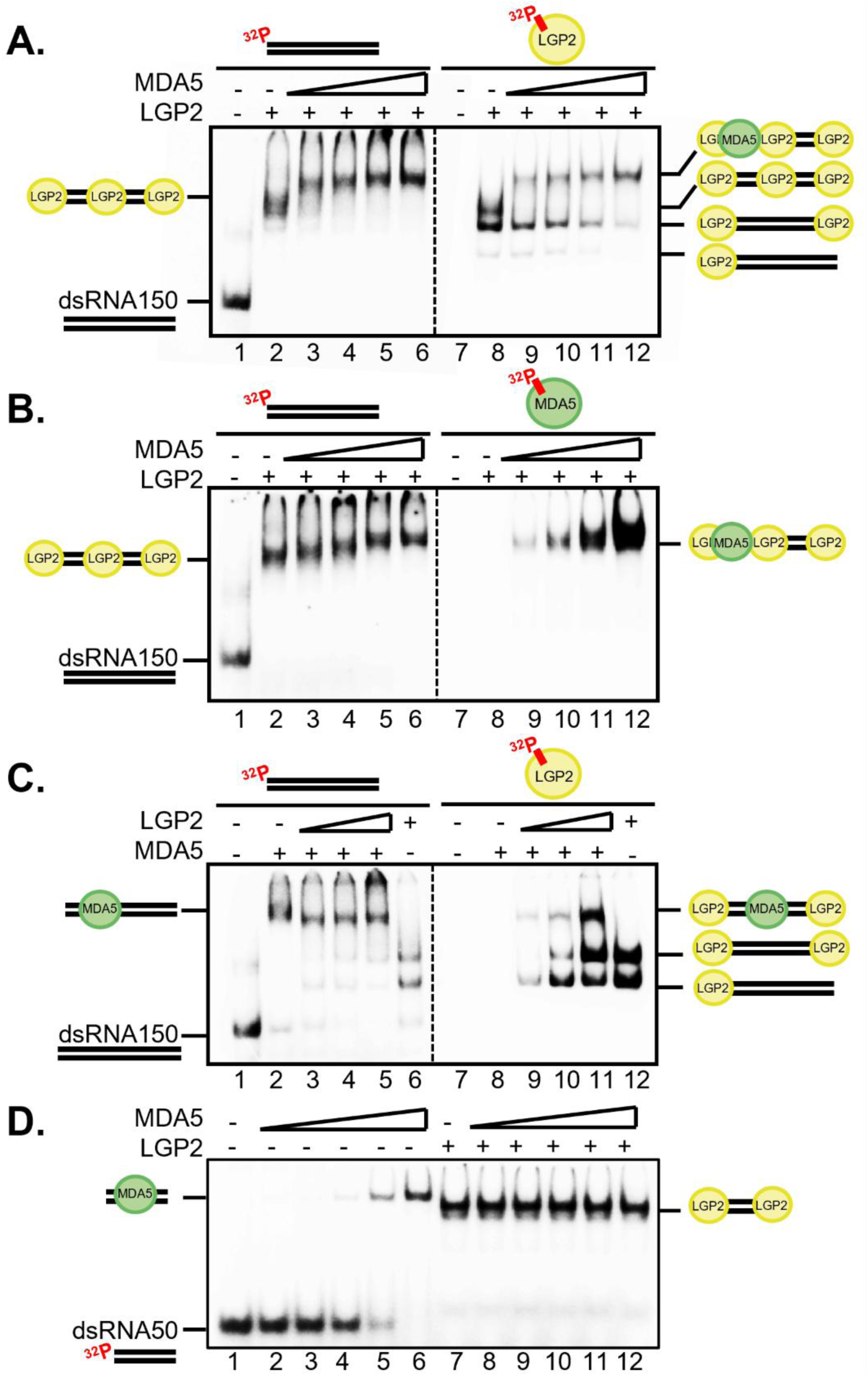
MDA5 and LGP2 form a stable heteromeric complex on synthetic dsRNA *in vitro.* **A**. EMSA performed using 20 nM dsRNA150-^32^P where 300 nM LGP2 was pre-loaded onto the dsRNA substrate (Lane 2). Increasing amounts of MDA5 (150, 200, 250, 300 nM) were added to the pre-existing LGP2-dsRNA150-^32^P complex (Lanes 3-6). Under the same EMSA conditions, dsRNA150 was not labeled and instead LGP2 was ^32^P-labeled to probe its presence in the heteromeric complex (Lanes 7-12). **B**. EMSA performed using 20 nM dsRNA150-^32^P where 300 nM LGP2 was pre-loaded onto the dsRNA substrate (Lane 2). Increasing amounts of MDA5 (150, 200, 250, 300 nM) were added to the pre-existing LGP2-dsRNA150-^32^P complex (Lanes 3-6). Under the same EMSA conditions, dsRNA150 was not labeled and instead MDA5 was ^32^P-labeled to probe its presence in the heteromeric complex (Lanes 7-12). **C**. EMSA performed using 20 nM dsRNA150-^32^P where 300 nM MDA5 was pre-loaded onto the dsRNA substrate (Lane 2). Increasing amounts of LGP2 (100, 200, 300 nM) were added to the pre-existing MDA5-dsRNA150-^32^P complex (Lanes 3-6). Under the same EMSA conditions, dsRNA150 was not labeled and instead LGP2 was ^32^P-labeled to probe its presence in the heteromeric complex (Lanes 7-12). **D**. EMSA performed using 20 nM dsRNA50-^32^P where increasing amounts of MDA5 (100, 150, 200, 250, 300 nM) were added (Lanes 2-6). MDA5 was added to a pre-existing LGP2-dsRNA150-^32^P complex, where 300 nM LGP2 was loaded (Lane 7), to access heteromeric complex assembly (Lanes 8-12).

To assess whether the order of protein addition plays a role in complex assembly, we next pre-assembled the MDA5-dsRNA150 complex (**Fig. 2C**, Lanes 1,2) and then added LGP2. This resulted in a change in mobility of the super-shifted band (**Fig. 2C**, Lanes 3-5) suggesting that the filament composition differs from that observed when MDA5 binds to dsRNA150 alone (**Fig. 2C**, Lane 6). When this same experiment was visualized with LGP2-^32^P (**Fig. 2C**, Lanes 7-12), a super-shifted band appeared (**Fig. 2C**, Lanes 9-11) that is not present when LGP2-^32^P binds by itself to dsRNA150 (**Fig. 2C**, Lane 12), establishing the addition of LGP2 to a pre-existing MDA5-dsRNA150 complex. Interestingly, the LGP2-MDA5-dsRNA150 complex migrated faster than the MDA5-dsRNA150 complex (**Fig. 2C**, Lanes 2-5), suggesting that LGP2 incorporation alters MDA5 filament architecture. Collectively, these experiments indicate that MDA5 and LGP2 coexist on synthetic dsRNA150 *in vitro,* regardless of which protein is loaded first, resulting in the formation of a structurally distinct heteromeric complex

To assess the dsRNA length dependence of LGP2-MDA5-dsRNA complex assembly *in vitro*, we then examined complex formation on a shorter radiolabeled dsRNA of 50 bp (dsRNA50-^32^P). MDA5 efficiently bound dsRNA50-^32^P, as a shift of band migration appeared with increasing amounts of MDA5 (**Fig. 2D**, Lanes 1-6). However, when LGP2 is pre-loaded onto dsRNA50-^32^P, with two molecules bound (**Fig. S2C**), the band migration remained unchanged when increasing amounts of MDA5 were added to the LGP2-dsRNA50 complex (**Fig. 2D**, Lanes 7-12). These findings indicate that LGP2 and MDA5 do not form a stable heteromeric complex on dsRNA50. We next examined heteromeric complex assembly on a longer synthetic dsRNA substrate, dsRNA300. As expected, both LGP2 and MDA5 efficiently bound dsRNA300 individually (**Fig. S3A,B**). To assess heteromeric complex assembly, we first preloaded LGP2 onto dsRNA300 to form a stable LGP2-dsRNA300 complex (**Fig. S3C**, Lanes 1,2) and then added MDA5. Upon addition of MDA5, we observed a super-shifted band, representing formation of a stable LGP2-MDA5-dsRNA300 complex (**Fig. S3C**, Lanes 3-5). Therefore, these experiments demonstrate that LGP2 and MDA5 form stable complexes on longer dsRNAs but not on shorter dsRNAs, indicating that heteromeric complex assembly is dependent on dsRNA length.

As we were attempting to visualize stable complexes, EMSAs were conducted without ATP to prevent rapid dissociation and recycling of MDA5 and LGP2 on dsRNA, which was reported in other studies and observed in our case well (12, 23). To assess whether ATP binding affects LGP2-MDA5-dsRNA complex assembly *in vitro*, we examined dsRNA binding and heteromeric complex formation in the presence of the non-hydrolysable ATP analog, adenosine-5’-[(β,γ)-imido]triphosphate (AMP-PNP). Under this condition, both LGP2 and MDA5 formed stable individual complexes with dsRNA300 (**Fig. S4A,B**), comparable to their behavior in the absence of nucleotide. We then conducted experiments on LGP2 and MDA5 binding together to dsRNA, in the presence and absence of AMP-PNP, to assess heteromeric complex assembly. We first preloaded LGP2 onto dsRNA300 to form a stable LGP2-dsRNA300 complex (**Fig. S4C**, Lanes 1,2 and 6,7) and then added MDA5. Upon addition of MDA5, we observed a super-shifted band, representing formation of a stable LGP2-MDA5-dsRNA300 complex in both the absence and presence of AMP-PNP (**Fig. S4C**, Lanes 3-5 and 8-10). These experiments demonstrate that ATP binding does not impact heteromeric complex assembly.

### LGP2 lowers the threshold for MDA5 filament nucleation

Although the experiments described above provided insight into MDA5 filament assembly in the presence of LGP2, they did not define the stoichiometry, compositional heterogeneity or distribution of these filament species. Moreover, given that LGP2 enhances MDA5 signaling, we also sought to determine whether LGP2 regulates filament stability. To address these questions directly, we employed mass photometry experiments in the absence and presence of LGP2. Mass photometry measures the molecular mass of individual biomolecules in solution through interferometric light scattering (42)(Refeyn Ltd). Because heterogeneous protein-nucleic acid complexes contain multiple molecular species, mass photometry resolves these species as discrete peaks in the observed mass distribution, with peak counts reflecting the abundance of each species (42, 43). As a result, each peak represents a distinct complex with a defined molecular mass, providing a direct readout of complex stoichiometry and molecular diversity within heterogeneous macromolecular assemblies (42, 43).

To initiate this study, we first measured the molecular mass of all components individually (**Fig. S5**). As expected, we found that LGP2 and MDA5 exhibited similar molecular masses (∼81 kDa and ∼86 kDa, respectively) and each yielded a singular peak (**Fig. S5A,B**). Moreover, both dsRNA150 and dsRNA300 yielded a single peak at the expected molecular masses (∼83 kDa and 152 kDa, respectively), indicating proper assembly of the dsRNA substrates (**Fig. S5C,D**). Having validated the integrity of each individual component, we next assembled complexes containing these components for mass photometry analysis.

We then set out to examine complexes of dsRNA150 and LGP2 alone. To monitor the stoichiometry of individual LGP2-dsRNA150 complexes, we acquired mass photometry measurements of complexes containing 15 nM LGP2 and 1 nM dsRNA150 (**Fig. 3A).** Because LGP2 was in large excess, we observed a peak corresponding to free LGP2 protein (∼79 kDa, 38%), along with complexes containing one (∼154 kDa, 39%), two (∼232 kDa, 17%), or three (∼311 kDa, 6%) molecules of LGP2 bound to dsRNA150 (**Fig. 3A**). The predominant LGP2-dsRNA150 species corresponds to a single LGP2 molecule bound to dsRNA150, with a substantial population of complexes containing two LGP2 molecules, consistent with the preferential binding of LGP2 to dsRNA termini (25–27). We also observed an internally bound LPG2 in a low abundance of 3LGP2-dsRNA150 complexes, consistent with previous reports that LGP2 exhibits reduced affinity for dsRNA stems (25, 29). Due to this reduced affinity, we suspect internally bound LGP2 molecules likely dissociated over the course of the experiment, therefore producing a lower abundance. These data demonstrate the formation of defined LGP2-dsRNA150 complexes and establish the expected stoichiometries for investigating how LGP2 regulates MDA5 filament assembly on dsRNA150.

**Figure 3.**
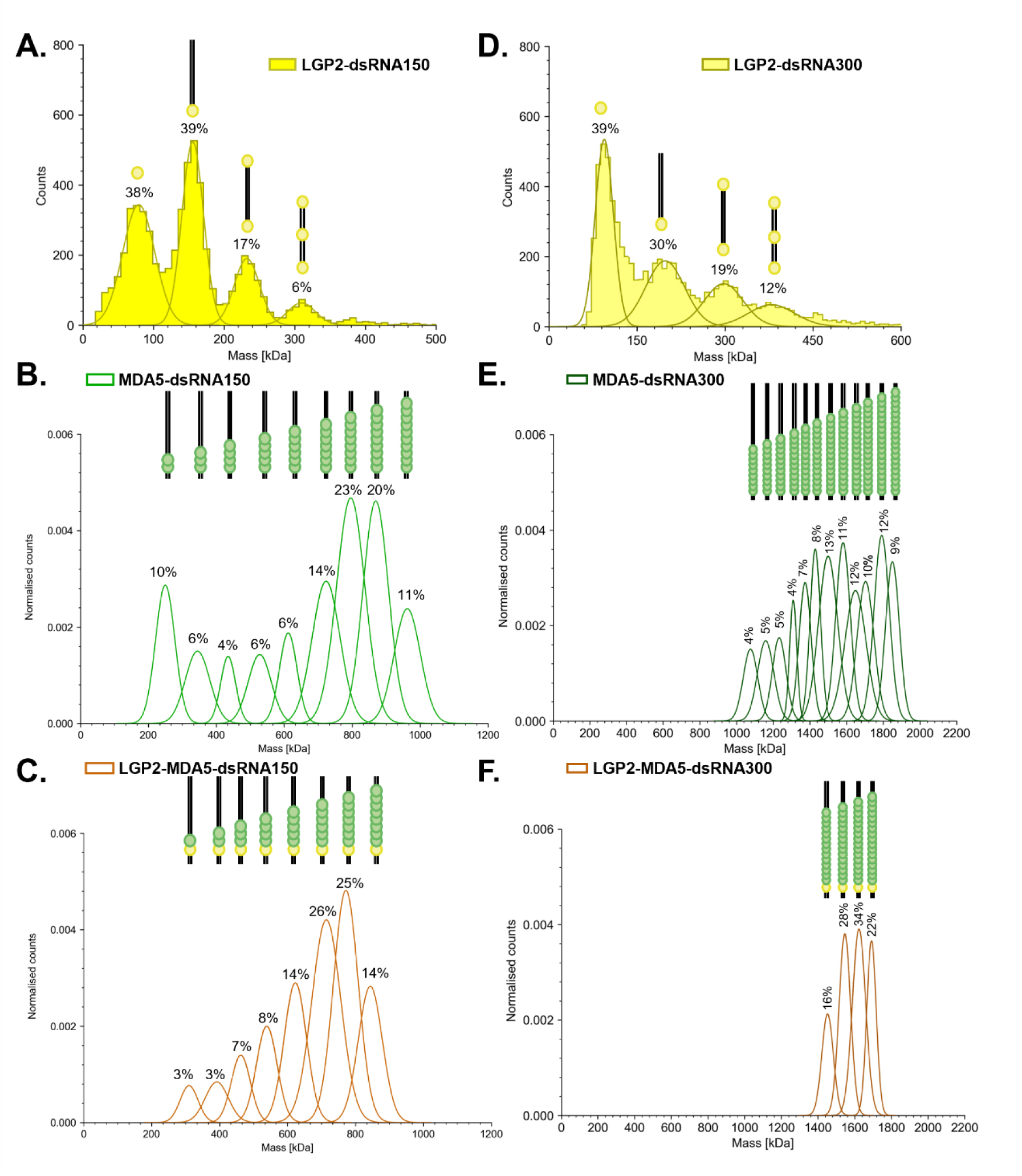
LGP2 allows for fewer molecules of MDA5 to nucleate stable filaments. **A.** Mass photometry measurement for the LGP2-dsRNA150 complex following manual 20-fold droplet dilution resulting in the final concentration of 15 nM LGP2 and 1 nM dsRNA150. Molecular weights correspond to free LGP2 (∼79 kDa), LGP2-dsRNA150 (∼156 kDa), 2LGP2-dsRNA150 (∼232 kDa), and 3LGP2-dsRNA150 (∼310 kDa), as depicted. Gaussian fitting is shown with quantification of areas under the curve to evaluate stoichiometric abundance. **B, C.** Mass photometry measurements for (**C**) MDA5-dsRNA150 and (**D**) LGP2-MDA5-dsRNA150 complexes utilizing the MassFluidix HC microfluidics system (Refeyn Ltd). Unbound protein peak at ∼86 kDa is not shown. Gaussian fitting is shown with quantification of areas under the curve to evaluate filament stoichiometric abundance between ∼150-1000 kDa. Histograms are shown in **Fig. S6A,B**. Schematic representation of predicted stoichiometric ratios of MDA5-dsRNA150 and LGP2-MDA5-dsRNA150 are shown. **D.** Mass photometry measurement for the LGP2-dsRNA300 complex following manual 20-fold droplet dilution resulting in the final concentration of 30 nM LGP2 and 1 nM dsRNA300. Molecular weights correspond to free LGP2 (∼94 kDa), LGP2-dsRNA300 (∼198 kDa), 2LGP2-dsRNA150 (∼298 kDa), and 3LGP2-dsRNA150 (∼379 kDa), as depicted. Gaussian fitting is shown with quantification of areas under the curve to evaluate stoichiometric abundance. **E, F.** Mass photometry measurements for (**E**) MDA5-dsRNA300 and (**F**) LGP2-MDA5-dsRNA300 complexes utilizing the MassFluidix HC microfluidics system (Refeyn Ltd). Unbound protein peak at ∼86 kDa is not shown. Gaussian fitting is shown with quantification of areas under the curve to evaluate filament stoichiometric abundance between ∼150-2000 kDa. Histograms are shown in **Fig. S8A,B**. Schematic representation of predicted stoichiometric ratios of MDA5-dsRNA300 and LGP2-MDA5-dsRNA300 are shown.

We next examined the composition and stoichiometry of MDA5-dsRNA150 filaments. As MDA5 assembles cooperatively along dsRNA to form filaments (23), each peak in the mass distribution is expected to represent a filament population containing a defined number of MDA5 molecules assembled along the dsRNA. The mass distribution of the MDA5-dsRNA150 complex revealed nine peaks, each peak representing a filament containing increasing amounts of MDA5 molecules bound to dsRNA150 (**Fig. 3B** and **Fig. S6A**). The largest MDA5 filament peak was observed at ∼962 kDa, which corresponds to ten molecules of MDA5 bound to one molecule of dsRNA150 (11%, **Fig. 3B** and **Fig. S5A**), consistent with a footprint of 14-15 bp per MDA5 molecule (21). The most abundant filament species contained either eight (∼796 kDa, 23%) or nine (∼870 kDa, 20%) molecules of MDA5 bound to dsRNA150 (**Fig. 3B** and **Fig. S5A**). We also observed a large proportion of intermediate species corresponding to 2-7 molecules of MDA5 bound to dsRNA150 (46%, **Fig. 3B** and **Fig. S5A**). We hypothesize that these intermediate filament species arise from partial disassembly of large filaments during the mass photometry experiment.

Finally, we characterized the composition of mixed LGP2-MDA5-dsRNA150 filaments to determine how LGP2 regulates MDA5 filament assembly and composition. The mass distribution of the LGP2-MDA5-dsRNA150 complex was encompassed by six peaks, each representing a filament with a defined number of MDA5 molecules bound to LGP2-dsRNA150 (**Fig. 3C** and **Fig. S5B**). Overall, the mass distribution appeared more uniform in the presence of LGP2, with a reduced number of filament species and a more even distribution of their relative abundances. To determine the stoichiometry of the complex, we assigned one LGP2 molecule per dsRNA150 due to predominance of the LGP2-dsRNA150 species observed by mass photometry measurements (39%, **Fig. 3A**). Here, the largest filament species was observed at ∼844 kDa, which corresponds to one molecule of LGP2 and eight molecules of MDA5 bound to dsRNA150 (14%, **Fig. 3C** and **Fig. S5B**). The most abundant species contained one molecule of LGP2 and either six (26%, ∼714 kDa) or seven (25%, ∼772 kDa) molecules of MDA5 bound to dsRNA150 (**Fig. 3C** and **Fig. S5B**). Therefore, the distinct stoichiometry of the largest filament species observed in the presence of LGP2 indicates that LGP2 reduces maximum number of MDA5 molecules required for filament nucleation. We also observed a population of intermediate filament species corresponding to complexes containing 1-5 molecules of MDA5 bound to LGP2-dsRNA150 (35%, **Fig. 3C** and **Fig. S5B**), although these species were less abundant than those observed for MDA5-dsRNA150. The reduction in intermediate filament species observed in the presence of LGP2 may reflect decreased disassembly of large filaments, consistent with enhanced filament stability. However, this stabilization effect is less pronounced on dsRNA150, as this shorter dsRNA substrate is less favorable for MDA5 filament assembly. Together, these findings indicate that LGP2 promotes both MDA5 filament nucleation and stabilization of full-length filaments.

To unambiguously determine the stoichiometry of the LGP2-MDA5-dsRNA150 complex, we fused a SUMO protein tag (small ubiquitin-like modifier, ∼13 kDa) to MDA5, increasing its molecular mass to distinguish it from LGP2 within the filament (**Fig. S7A**). When comparing the molecular masses of the largest LGP2-MDA5-dsRNA150 and LGP2-SUMO MDA5-dsRNA150 filament species, we identified peaks at ∼845 kDa and ∼957 kDa, respectively (**Fig. S7B**). This ∼112 kDa shift in molecular mass upon addition of the SUMO tag to MDA5 corresponds to roughly eight SUMO tags, confirming that the largest filament species contains eight MDA5 molecules and one LGP2 molecule (**Fig. S7C**). These findings provide direct experimental evidence that, in solution, each MDA5 filament is associated with a single LGP2 molecule.

Because MDA5 preferentially forms filaments on longer dsRNA substrates (40), we carried out a parallel analysis using a longer dsRNA (dsRNA300) to assess whether LGP2 regulates MDA5 filament assembly in a manner that differs from shorter dsRNAs. To this end, we first performed mass photometry experiments to determine the stoichiometry of individual LGP2-dsRNA300 complexes containing 30 nM LGP2 and 1 nM dsRNA300 (**Fig. 3D**). Despite the increased dsRNA length, we observed a similar relative binding stoichiometry on dsRNA300 as observed on dsRNA150. Mass photometry measurements identified LGP2-dsRNA300 complexes containing one (∼198 kDa, 30%), two (∼298 kDa, 19%), or three (∼379 kDa, 12%) molecules of LGP2 bound to dsRNA300 (**Fig. 3D**). Strikingly, the predominant species contained one LGP2 molecule bound to dsRNA300, with a substantial population of complexes containing two LGP2 molecules. This distribution is consistent with the end-binding preference of LGP2 (25–27), as increasing dsRNA length does not alter LGP2 binding stoichiometry or promote more than one internal dsRNA binding event.

To monitor MDA5 filament formation by mass photometry, we first characterized filaments of MDA5 alone on dsRNA300. The mass profiles revealed twelve distinct peaks for the MDA5-dsRNA300 complex, each exhibiting a relative abundance of 4-12% (**Fig. 3E** and **Fig. S8A**). Similar to dsRNA150, many short filament species were observed, potentially generated through partial disassembly of larger filaments during the experiment, resulting in a broad diversity of distinct filament populations. The largest species contained 20 molecules of MDA5 bound to dsRNA300 (9%, ∼1850 kDa, **Fig. 3E** and **Fig. S8A**), consistent with a footprint of 14-15 bp per MDA5 molecule (21). These data are consistent with the results obtained with dsRNA150 and demonstrate that dsRNA300 accommodates twice the number of MDA5 molecules, as expected for a dsRNA substrate that is twice as long.

We next monitored the characteristics of mixed LGP2-MDA5-dsRNA300 filaments to determine how LGP2 regulates MDA5 filament assembly and stability on this longer dsRNA substrate. In contrast to MDA5 alone, we observed only four distinct types of filaments, and a more uniform abundance distribution was observed for the LGP2-MDA5-dsRNA300 complexes (16-34%, **Fig. 3F** and **Fig. S8B**). In these complexes, which contained one molecule of LGP2 bound, the largest species contained 17 molecules of MDA5 (22%, ∼1692 kDa, **Fig. 3F** and **Fig. S8B**). Consistent with the dsRNA150 data, these results indicate that LGP2 reduces the number of MDA5 molecules required for maximal filament nucleation. Additionally, LGP2 drastically reduces the number of low-molecular weight filament species and promotes the persistence of relatively long filaments, suggesting that LGP2 enhances the stability of long filaments. Importantly, the stabilizing effect of LGP2 is more pronounced with dsRNA300.

### LGP2 reduces disassembly of MDA5 filaments

While the mass photometry experiments conducted thus far enabled determination of protein stoichiometry and relative filament size distribution, they did not provide a direct quantitative measure of filament stability. To address this, we implemented a variation in the mass photometry experiment, much like a pulse chase experiment (44), which enabled us to estimate the dissociation rate constants (k_off_^app^) of the filament assemblies. Complexes were first allowed to reach equilibrium, and then filament disassembly was initiated by dilution and monitored over a five-minute observation period. Filament populations (∼239-2000 kDa) were quantified at each time point as the number of species within a given mass range. Specifically, filaments were classified as intermediate (∼239-800 kDa) or large (∼800-2000 kDa), with intermediate filaments interpreted as products of large filament disassembly. The time-dependent decrease in the abundance of large filament species was then used to quantify filament disassembly kinetics and assess filament stability.

First, we examined the mass distributions of MDA5-dsRNA300 complexes over the five-minute observation period following dilution. At the initial time point, MDA5-dsRNA300 filaments spanned a broad mass range containing both intermediate and large filament species (**Fig. S9A**). Over the course of the experiment, we observed a substantial decrease in the abundance of intact filaments (**Fig. S9A**), consistent with progressive filament disassembly. We next examined LGP2-MDA5-dsRNA300 filaments under the same conditions and observed pronounced differences in the mass distribution. At the initial time point, LGP2-MDA5-dsRNA300 filaments exhibited a more uniform distribution with a greater proportion of larger filaments relative to intermediate filaments (**Fig. S9B**). After five minutes, LGP2-MDA5-dsRNA300 complexes retained a large proportion of large filament species and a higher overall abundance of intact filaments than that observed with MDA5-dsRNA300 complexes (**Fig. S9C**), consistent with enhanced filament stability in the presence of LGP2.

We then sought to measure filament dissociation kinetics and thereby quantitatively compare MDA5 filament stability in the absence and presence of LGP2. The fraction of intact large filament species (≥800 kDa) was quantified over time by mass photometry and fit to a one-phase exponential decay model for MDA5-dsRNA300 (dark green line, R^2^= 0.99) and LGP2-MDA5-dsRNA300 (dark orange line, R^2^ = 0.99) complexes (**Fig. S9D**). Using this approach, we determined apparent dissociation rate constants (k_off_^app^) from the resulting decay curves, yielding values of 0.57±0.12 min^−1^ and 0.31±0.10 min^−1^ for MDA5-dsRNA300 and LGP2-MDA5-dsRNA300 complexes, respectively (**Fig. S9D**). The approximate two-fold reduction in the apparent dissociation rate in the presence of LGP2 indicates that LGP2 increases MDA5 filament stability by reducing the rate of filament disassembly.

### MDA5 and LGP2 directly interact in a dsRNA-dependent manner

Given that MDA5 and LGP2 appear to form a stable complex, we set out to determine whether specific molecular interactions between LGP2 and MDA5 enhance filament assembly and stability. To screen for molecular interactions between MDA5 and LGP2 *in vitro*, we performed protein-protein crosslinking experiments using the cross-linker disuccinimidyl glutarate (DSG), which covalently links lysine residues that are positioned within 7.7 Å apart. This approach enabled us to capture protein-protein complexes and visualize them by SDS-PAGE analysis. We first incubated radiolabeled MDA5 (MDA5-^32^P) with LGP2 to probe the existence of protein-protein interactions in the absence of dsRNA. Upon addition of DSG, we did not observe a distinct crosslinked product at the expected molecular weight of the covalently linked LGP2-MDA5 complex (∼160 kDa), indicating that LGP2 and MDA5 do not interact under these conditions (**Fig. 4A**, Lanes 1,2). We next examined whether dsRNA promotes LGP2-MDA5 interactions by introducing high molecular weight (HMW) poly(I:C) as a MDA5 filament substrate. Although LGP2-MDA5 interactions were not detected at very low concentrations of poly(I:C) (**Fig. 4A**, Lanes 2-4), higher poly(I:C) concentrations promoted formation of an LGP2-MDA5 crosslinked product (**Fig. 4A**, Lanes 5-9). With increasing amounts of poly(I:C), additional crosslinked products were observed, corresponding to a covalently linked MDA5-MDA5 complex based on its molecular weight (∼170 kDa, **Fig. 4A**, Lanes 5-9), which is consistent with MDA5 oligomerization on dsRNA. Together, these results indicate that LGP2-MDA5 interactions occur in a dsRNA-dependent manner and are associated with MDA5 oligomerization on dsRNA.

**Figure 4.**
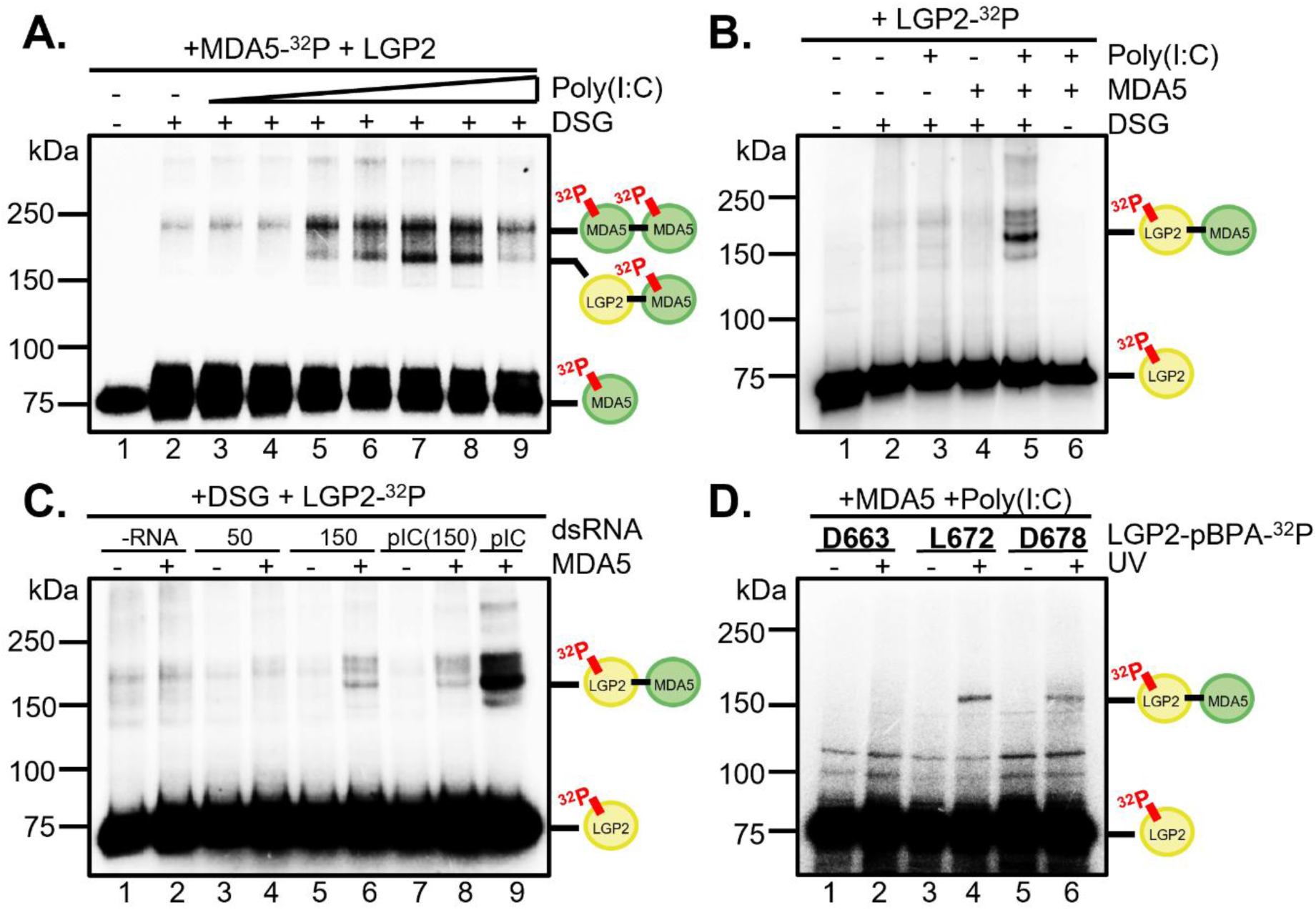
MDA5 and LGP2 directly interact in a dsRNA-dependent manner. **A.** DSG crosslinking performed using 200 nM MDA5-^32^P with increasing amounts of HMW poly(I:C) (0.05, 0.1, 0.5, 1.0, 5.0, 10, 50 ug/ml, Lanes 3-9) in the presence of 200 nM LGP2. **B.** DSG crosslinking performed using 200 nM LGP2-^32^P in the absence and presence of 200 nM MDA5 and/or 5 ug/ml HMW poly(I:C). **C.** DSG crosslinking performed using 200 nM LGP2-^32^P in the absence and presence of 200 nM MDA5 and 5 ug/ml dsRNA substrates. Crosslinking in the presence of 5 ug/ml HMW pIC was used as a positive control (Lane 9). **D.** The site-specific photo-crosslinking probe, parabenzoyl phenylalanine (pBPA), was incorporated into the CTT of LGP2-^32^P at residues D663, L672, or D678. Crosslinking reactions contained 100 nM LGP2-pBPA-^32^P, 100 nM MDA5, and 1 ug/ml HMW poly(I:C) and subjected to UV irradiation (360 nm) for 20 minutes, where indicated. Bands around ∼100 kDa represent non-specifically radiolabeled contaminants.

To unambiguously determine whether the crosslinked species contains LGP2, we performed a complementary crosslinking experiment in which LGP2 was radiolabeled (LGP2-^32^P) and MDA5 was left unlabeled. We first incubated LGP2-^32^P alone with DSG in the absence or presence of poly(I:C) and did not detect any crosslinked products (**Fig. 4B**, Lanes 2,3), indicating that LGP2 does not self-associate. In the absence of poly(I:C), addition of MDA5 to the reaction containing LGP2-^32^P did not result in detectable crosslinked products (**Fig. 4B**, Lane 4). However, abundant crosslinked products were detected when LGP2-^32^P, MDA5, and 5 ug/ml poly(I:C) were incubated together in the presence of DSG (**Fig. 4B**, Lane 5). The molecular weight of this crosslinked product is consistent with a covalently linked LGP2-MDA5 complex (∼160 kDa, **Fig. 4B**, Lane 5), providing additional evidence that MDA5 and LGP2 interact only in the presence of dsRNA, and that both proteins are present in the complex.

The previous experiments were conducted with the unnatural polymer poly(I:C), so we set out to determine whether LGP2 and MDA5 crosslinks could be observed on mixed-sequence dsRNA substrates that support heteromeric complex assembly *in vitro*. In the initial experiments, LGP2-^32^P was incubated with DSG in the absence and presence of MDA5 and dsRNA (**Fig. 4C**). As expected, crosslinked products were not observed when MDA5 was absent (**Fig. 4C**, Lanes 1,3,5,7). Crosslinked products were not observed in the absence of dsRNA and in the presence of the relatively short dsRNA50 duplex (**Fig. 4C**, Lanes 3,4). However, LGP2-MDA5 crosslinks were observed in the presence of intermediate-length dsRNAs such as dsRNA150 and a poly(I:C) substrate 150 bp in length (pIC(150)) (∼160 kDa, **Fig. 4C**, Lanes 6,8). These experiments demonstrate that an interaction interface between LGP2 and MDA5 only forms on longer dsRNA substrates that support heteromeric complex formation.

### The C-terminal tail of LGP2 mediates LGP2-MDA5 interactions

Collectively, these findings prompted us to investigate the LGP2-mediated molecular interactions that drive MDA5 filament assembly and stability. Recent cryo-EM studies elucidated three LGP2-MDA5 interaction interfaces, one including the C-terminal tail of LGP2 (CTT, residues 663-678) (31). The reported LGP2-MDA5 interface is consistent with previous structural predictions and functional mutagenesis studies, all of which suggest a role for the CTT of LGP2 in mediating LGP2-MDA5 interactions (29). These findings prompted us to further examine the role of the CTT of LGP2 in heteromeric complex assembly and function. We first employed a site-specific protein-protein crosslinking approach to validate a CTT-mediated LGP2-MDA5 functional interaction interface. To this end, we incorporated the photo-reactive amino acid p-benzoyl-L-phenyl alanine (pBPA) into various sites of the CTT of LGP2 using an *E. coli* expression system (45). Based on sequence conservation across mammalian species, we substituted pBPA into three conserved residues of interest, D663, L672, and D678 (**Fig. S10A**). For pBPA crosslinking experiments, UV activation induces covalent crosslinking of pBPA residues that come into contact with a neighboring amino acid. Following SDS PAGE analysis, the presence of a higher molecular weight crosslinked product indicates a direct interaction between MDA5 and LGP2 at the pBPA-substituted residue. To initiate crosslinking, LGP2-^32^P with pBPA incorporated into the CTT was incubated with MDA5 and HMW poly(I:C) and subjected to UV exposure (**Fig. S10B**). Following UV exposure, one distinct band corresponding to a covalently linked LGP2-MDA5 complex (∼160 kDa) appeared when pBPA was incorporated into residues L672 and D678 (**Fig. 4D**, Lanes 4,6), but not D663 (**Fig. 4D**, Lane 2). These experiments demonstrate that the terminal end of the CTT is directly involved in the LGP2-MDA5 interface and prompted us to further characterize this interaction and determine its role in regulating MDA5 filament assembly.

### Deletion of the C-terminal tail of LGP2 disrupts the LGP2-MDA5 interface

We next investigated whether the CTT of LGP2 mediates recruitment of MDA5 to dsRNA and promotes heteromeric complex formation. To test the functional importance of this interaction interface, we generated an LGP2 deletion mutant lacking CTT residues 663-678 (“ΔCTT LGP2”). We found that deletion of the CTT did not impair ATPase activity (**Fig. S11A**) or dsRNA binding activity (**Fig. S11B**), indicating that the mutant enzyme retains its core functions. We then used the ΔCTT LGP2 mutant to test whether the LGP2 CTT-MDA5 interaction interface is required for heteromeric complex formation by EMSA. We first loaded ΔCTT LGP2 onto dsRNA150-^32^P and observed a shift in band migration representing a ΔCTT LGP2-dsRNA150 complex (**Fig. 5A**, Lanes 1,2). Upon addition of MDA5 to the pre-formed ΔCTT LGP2-dsRNA150 complex, a “super-shifted” band did not appear (**Fig. 5A**, Lanes 3-6), indicating MDA5 cannot bind. Instead, we observed a slower migrating band that likely corresponds to MDA5 competitively binding to dsRNA150 alone (**Fig. 5A**, Lane 6). To further examine whether MDA5 is part of the heteromeric complex, we radiolabeled MDA5 with ^32^P (MDA5-^32^P) and left dsRNA150 unlabeled (**Fig. 5A**, Lanes 9-12). When monitoring MDA5-^32^P, we observed no discrete bands with increasing amounts of MDA5-^32^P were added to the pre-formed ΔCTT LGP2-dsRNA150 complex **(Fig. 5A**, Lanes 9-12). These results indicate that ΔCTT LGP2 and MDA5 cannot form a stable heteromeric complex on dsRNA150, as ΔCTT LGP2 severely impairs MDA5 recruitment to dsRNA150.

**Figure 5.**
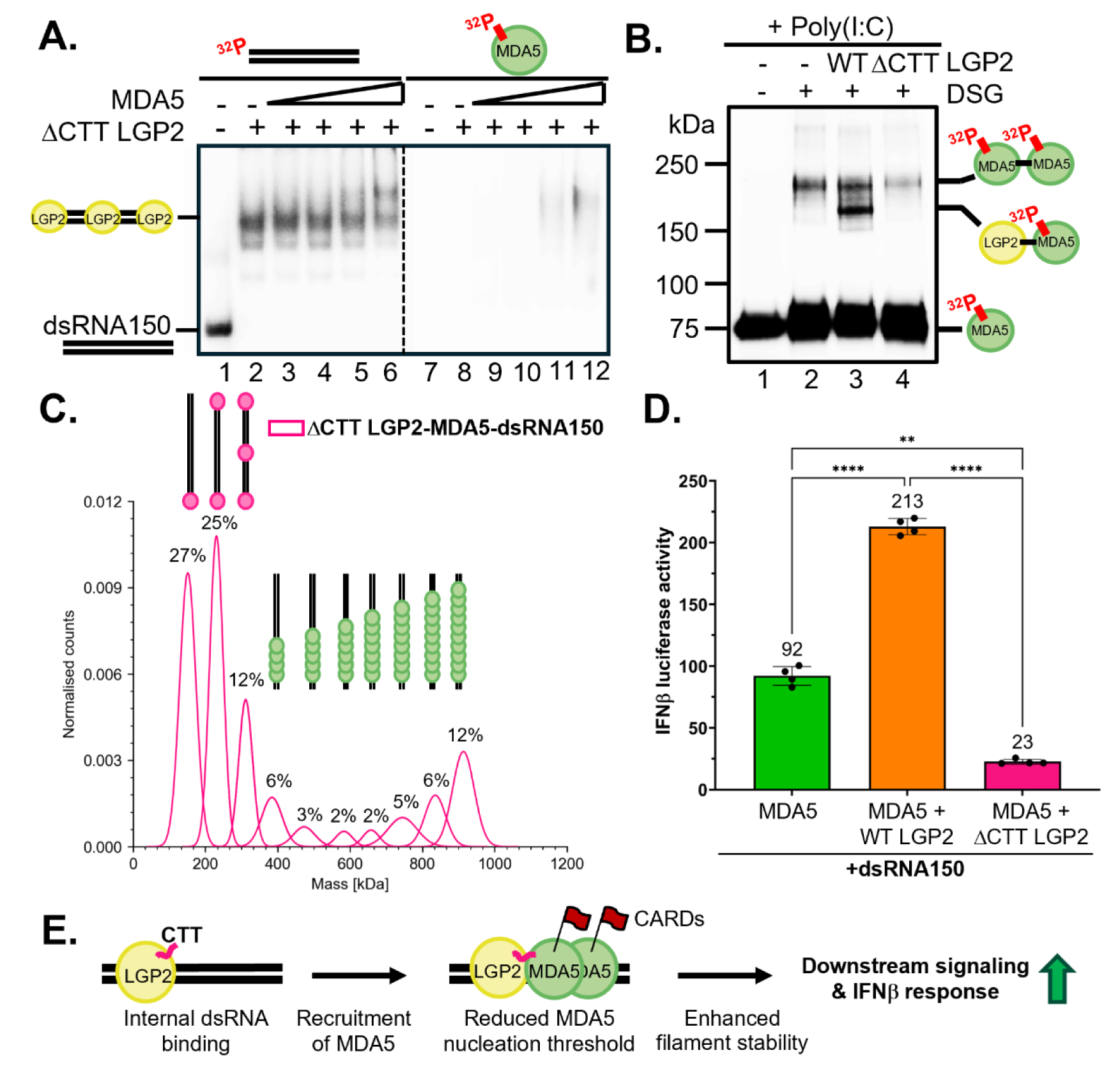
Deletion of the C-terminal tail of LGP2 disrupts the LGP2-MDA5 interface. **A.** EMSA performed using 20 nM dsRNA150-^32^P where 300 nM ΔCTT LGP2 was pre-loaded to the dsRNA substrate (Lane 2). Increasing amounts of MDA5 (150, 200, 250, 300 nM) were added to the pre-existing ΔCTT LGP2-dsRNA150-^32^P complex (Lanes 3-6). Under the same EMSA conditions, dsRNA150 was not labeled and instead MDA5 was ^32^P-labeled to probe its presence in the heteromeric complex (Lanes 7-12). **B.** DSG crosslinking performed using 200 nM MDA5-^32^P in the presence of 5 ug/ml HMW poly(I:C) with 200 nM WT (Lane 3) or ΔCTT LGP2 (Lane 4) added to the reaction. **C.** Mass photometry measurements for the ΔCTT LGP2-MDA5-dsRNA150 (pink) complex utilizing the MassFluidix HC microfluidics system (Refeyn Ltd). Unbound protein peak at ∼86 kDa is not shown. Gaussian fitting is shown with quantification of areas under the curve to evaluate filament stoichiometric abundance. Histograms are shown in **Fig. S12C**. Schematic representation of predicted stoichiometric ratios of ΔCTT LGP2-dsRNA150 and MDA5-dsRNA150 are shown. **D.** IFN induction by co-transfection of MDA5 (1 ng) and WT or ΔCTT LGP2 (5 ng) plasmids stimulated with 1 ug dsRNA150 in HEK293T cells, where indicated. Data are represented as mean±SD (n=3 biological replicates). ****p<0.0001 and **p=0.0029 by one-way ANOVA. **E.** Working model of LGP2-mediated enhancement of MDA5 signaling. LGP2 first binds dsRNA due to its faster on-rate and recruits MDA5 through direct LGP2 CTT-MDA5 interactions. The heteromeric complex is stabilized by these protein-protein interactions and allows for increased downstream signaling and IFNβ response through enhanced MDA5 filament stability.

These results prompted us to assess whether any other interactions were taking place once the CTT of LGP2 was removed by DSG protein-protein crosslinking. In the presence of DSG, 5 ug/ml HMW poly(I:C), and MDA5-^32^P, MDA5 crosslinked with itself due to oligomerization on dsRNA (∼170 kDa, **Fig. 5B**, Lane 2). Upon addition of wild type (WT) LGP2 to the reaction, a covalently linked LGP2-MDA5 complex appeared (∼160 kDa, **Fig. 5B**, Lane 3), representing the presence of functional interactions between the two proteins. However, when ΔCTT LGP2 was added to the reaction instead, we did not observe formation of an LGP2-MDA5 crosslinked product (**Fig. 5B**, Lane 4), indicating that the CTT of LGP2 is required for LGP2-MDA5 interactions. Moreover, the band corresponding to MDA5-MDA5 oligomerization decreased significantly in the presence of ΔCTT LGP2 (**Fig. 5B**, Lane 4) compared with reactions containing WT LGP2 (**Fig. 5B**, Lane 3). Taken together, this experiment indicates that deletion of the CTT of LGP2 eliminates functional interactions with MDA5 and perturbs MDA5 oligomerization on dsRNA.

Given these findings, we next examined MDA5-dsRNA150 filament stoichiometry and composition in the presence of ΔCTT LGP2 by mass photometry. We first performed mass photometry experiments to determine the stoichiometry of individual ΔCTT LGP2-dsRNA150 complexes assembled from 15 nM ΔCTT LGP2 and 1 nM dsRNA150 (**Fig. S12A,B**). Mass photometry revealed a population of free ΔCTT LGP2 protein (∼88 kDa, 43%) and complexes containing one (∼153 kDa, 37%), two (∼227 kDa, 16%), or three (∼304 kDa, 4%) ΔCTT LGP2 molecules bound to dsRNA150 (**Fig. S12B**). This stoichiometry distribution closely resembled that observed for WT LGP2, indicating that deletion of the CTT does not alter the dsRNA-binding behavior of LGP2. We next examined ΔCTT LGP2-MDA5-dsRNA150 complexes by mass photometry. The mass profile resolved ten distinct peaks spanning a broad range of molecular masses (**Fig. 5C** and **Fig. S12C**). The majority of peaks (64%) correspond to ΔCTT LGP2 binding to dsRNA150, with peaks detected at approximately ∼152 kDa (27%), ∼231 kDa (25%), and ∼311 kDa (12%, **Fig. 5C** and **Fig. S12C**). The remaining peaks (36%) correspond to higher molecular weight filament species spanning approximately ∼384-914 kDa, with the most abundant species having a molecular mass of ∼914 kDa (12%, **Fig. 5C** and **Fig. S12C**). Notably, the molecular mass distribution of these higher molecular weight filament species more closely resembled MDA5-dsRNA150 filaments than WT LGP2-MDA5-dsRNA150 filaments (**Fig. 3B,C**). Therefore, the predominant filament peak at ∼914 kDa corresponds to a complex containing approximately ten molecules of MDA5 bound to dsRNA150 (**Fig. 5C** and **Fig. S12C**). We also observed a high abundance of intermediate filament species containing approximately 4-9 molecules of MDA5 bound to dsRNA150 (24%, **Fig. 5C** and **Fig. S12C**), consistent with destabilized MDA5 filaments and partial filament disassembly. Therefore, these results demonstrate that these filament species consist primarily of MDA5 without ΔCTT LGP2 bound to dsRNA. Taken together, these mass photometry data provide evidence that deletion of the CTT of LGP2 impairs LGP2-mediated MDA5 filament assembly.

Finally, to determine the functional consequences of ΔCTT LGP2 *in cellulo*, we monitored its ability to regulate MDA5 activity in the well-established cell-based reporter system for interferon signaling. Following transfection with dsRNA150, interferon signaling was observed as MDA5 is activated (green bar, **Fig. 5D**). Upon addition of WT LGP2, we observed a significant enhancement in MDA5-mediated interferon signaling (orange bar, ****p<0.0001, **Fig. 5D**). However, in the presence of ΔCTT LGP2 (pink bar, **Fig. 5D**), there was a significant decrease in MDA5-mediated interferon signaling relative to both WT LGP2 (****p<0.0001, **Fig. 5D**) and MDA5 alone (**p=0.0029, **Fig. 5D**). This result suggests that deletion of the CTT of LGP2 disrupts LGP2-MDA5 complex formation and reduces overall MDA5 signaling due to competitive dsRNA binding. Collectively, these findings demonstrate that the CTT of LGP2 serves as a key LGP2-MDA5 interaction interface that enhances MDA5-mediated immune signaling (**Fig. 5E**).

## Discussion

Since its original discovery in 2001(4), the biological function of LGP2 has been particularly intriguing as it lacks the CARDs necessary for signaling and yet it has an overall structure and RNA-binding properties similar to those of MDA5 and RIG-I (6, 27, 36–38). Accumulating evidence has established that LGP2 expression influences the signaling efficiency of MDA5 and RIG-I, demonstrating that LGP2 plays an important role in the innate immune response (11–16). How LGP2 exerts this regulatory effect while lacking CARDs has remained a mystery until recently. Structural studies established that, like RIG-I (46), LGP2 is a selective dsRNA end-binder capable of recognizing viral dsRNA targets (25–27). This behavior suggests that LGP2 may function as an anchor on dsRNA, recruiting other proteins (such as MDA5) that possess CARDs capable of initiating downstream signaling. In this model, LGP2 may function analogously to CD14, which acts as a co-receptor for Toll-like receptor 4 (TLR4) by facilitating ligand recognition and transfer to the TLR4-MD-2 signaling complex to promote innate immune signaling (47, 48). Consistent with this model, previous biochemical and cryo-EM studies have shown that LGP2 can form complexes with MDA5 once LGP2 translocates away from dsRNA termini (12, 29, 31). However, whether LGP2 remains associated with MDA5-dsRNA signaling complexes after filament nucleation, and how it influences the organization of these complexes, has remained unclear.

Here, using quantitative biochemical and cell-based assays, we provide evidence supporting a model in which LGP2 serves as a specific and integral component of MDA5-dsRNA signaling complexes, contributing not only to MDA5 recruitment but also to the organization and function of these complexes. Although multiple studies established that LGP2 promotes MDA5 filament nucleation (12, 29–31, 33), the nature of signaling-competent heteromeric LGP2-MDA5 filaments has remained unclear. Features of the heteromeric complex have been difficult to resolve using conventional electron microscopy, as the similar architecture of LGP2 and MDA5 when bound to dsRNA limits the ability to unambiguously distinguish them and to identify their relative positions (12, 29–31, 33). Building on previous evidence for LGP2 incorporation into MDA5 filaments (30), our findings further support a broader role for LGP2 as an active functional component of the signaling complex.

We provide direct evidence that LGP2 reduces the number of MDA5 molecules required for stable filament formation on dsRNAs and that LGP2 association alters MDA5 filament architecture (**Fig. 3**). Because filament stoichiometry measured by mass photometry reflects filament length along dsRNA, these findings are consistent with previous electron microscopy studies reporting that LGP2 promotes the formation of shorter MDA5 filaments (12, 29, 31, 33). Although other studies have suggested that LGP2 induces conformational changes in MDA5 that promote its transition to a signaling-competent state (30), the molecular architecture of LGP2-MDA5 complexes within the filament remains an important area of study. The mass photometry experiments provided in this study reveal distinct molecular mass distributions of the heteromeric complex that are consistent with changes in MDA5 filament architecture when LGP2 is bound (**Fig. 3**). Further, the accompanying EMSA experiments show distinct migration patterns for LGP2-MDA5-dsRNA complexes relative to MDA5-dsRNA complexes (**Fig. 2**). Together, these findings support a model in which LGP2 promotes the formation of structurally distinct MDA5 filaments, representing a specific MDA5 filament isoform that facilitates a more robust immune response.

Given that stable MDA5-dsRNA complex formation is directly linked to signaling efficiency (40), quantifying MDA5 filament integrity in the absence and presence of LGP2 was essential. While previous studies examined ATP-dependent MDA5 filament disassembly, they did not quantitively characterize the effect of LGP2 on filament stability and turnover (30). Here, mass photometry enabled us to directly investigate the influence of LGP2 on filament size and the kinetics of filament disassembly. These studies revealed that LGP2 stabilizes MDA5 filaments, increasing their persistence and preventing disassembly (**Fig. S9**), thereby enhancing their ability to undertake a full cycle of signal induction. LGP2 may enhance filament stability by altering MDA5 organization, potentially by condensing the MDA5 spacing, in agreement with recent studies (33). Together, these findings establish a key role for LGP2 in maintaining stable MDA5 filaments, which helps explain the role of LGP2 in promoting efficient immune activation.

Notably, a recent cryo-EM study advanced our structural understanding of LGP2-mediated regulation of MDA5 signaling by identifying multiple LGP2-MDA5 interaction interfaces, including one involving the LGP2 CTT (31). However, in that work, the terminal six residues of the LGP2 CTT were not resolved and the functional role of the CTT LGP2-MDA5 interaction interface was not evaluated (31). Instead, the capping loop of LGP2 (residues 592-609) was highlighted as a primary LGP2-MDA5 interface (31), in contrast with previous hypotheses regarding the functional importance of the CTT (29). We establish here that the functional LGP2-MDA5 interface is broader, as it includes conserved residues in the CTT. While the capping loop may indeed contribute to the complex, it is important to note that the cryo-EM structures were obtained using high-affinity mutant LGP2 and MDA5 proteins (31), each of which contained gain-of-function (GOF) mutations known to artificially strengthen dsRNA association (32, 49, 50). Whether these substitutions alter the LGP2-MDA5 interface or preferentially stabilize a conformation also accessible to WT proteins remains to be determined. Nonetheless, the high-resolution structural findings support the formation of a defined heteromeric LGP2-MDA5 complex and identify the LGP2 CTT-MDA5 interface as a potentially critical interaction site (31). Together with our experimental and phylogenetic analyses (**Fig. 5** and **Fig. S10A**), these findings provide functional support for a model in which direct contacts between LGP2 and MDA5 coordinate their functional interplay to enhance MDA5-mediated antiviral signaling.

Consistent with a model in which LGP2 is an obligate cofactor of MDA5, LGP2 and MDA5 RNA expression levels are directly correlated in cells that depend heavily on innate immune signaling, such as dendritic cells (51)(Human Protein Atlas, proteinatlas.org). Additionally, LGP2 and MDA5 protein expression levels are directly correlated in tissues with high dendritic cell population, including skin, mucous membranes, spleen, and lymph nodes (51)(Human Protein Atlas, proteinatlas.org). These expression patterns support a mechanistic model in which LGP2 and MDA5 usually operate together, as a functional heteromeric complex during dsRNA sensing and signaling. Given the importance of MDA5 signaling in antiviral function and, when dysregulated, in autoimmunity, an understanding of the LGP2-MDA5 interface is particularly critical. Therefore, by further defining the role of LGP2 in MDA5 signaling, this work provides new insights into the basic mechanism by which MDA5 triggers the interferon response and opens the door to new therapeutic modalities that capitalize on the key role that LGP2 plays during the innate immune response.

## Materials and Methods

### Cloning, expression, and purification of proteins

N-terminal hexa-histidine-tagged human ΔCARDs MDA5 (residues 298-1025) and LGP2 were cloned into the pET-SUMO expression vector and mutations were introduced by the Quik-Change II XL Site-Directed Mutagenesis Kit (Agilent). Mutagenesis primers used in this study are listed in **Table S1**. Sequences were validated by whole-plasmid sequencing (Quintara). ΔCARDs MDA5 was expressed in Rosetta 2(DE3) *E. coli* cells (Novagen) and cells were grown in autoclaved LB media until the OD600 reached 0.6 and expression was induced by addition of 0.5 mM IPTG for 16 hours at 16°C. LGP2 was expressed in Rosetta 2(DE3) E. coli cells (Novagen) in filtered LB supplemented with 50 mM KH_2_PO_4_, 50 mM Na_2_HPO_4_, 25 mM (NH_4_)_2_SO_4_, 2 mM MgSO_4_, 0.5% glycerol, and 0.05% glucose, and expression was induced by addition of 0.25 mM IPTG when OD600 of the culture reached 6. Cells were harvested, resuspended in lysis buffer (50 mM NaH_2_PO_4_-Na_2_HPO_4_ pH 7.4, 300 mM NaCl, 10% glycerol, 10 mM imidazole, 5 mM 2-mercaptoethanol) supplemented with EDTA-free protease inhibitor cocktail (MedChemExpress), lysed by sonication, and clarified by centrifugation. The proteins were purified by affinity chromatography using Ni-NTA Agarose (Invitrogen), followed by affinity chromatography using a HiTrap™ Heparin HP column (Cytiva). The heparin column was equilibrated in buffer A (25 mM HEPES-NaOH pH 7.4, 150 mM NaCl, 5% glycerol, 5 mM 2-mercaptoethanol) and eluted by 0-90% linear gradient of buffer B (25 mM HEPES-NaOH pH 7.4, 1.5 M NaCl, 5% glycerol, 5 mM 2-mercaptoethanol). Peak fractions were collected and treated with in-house-prepared ULP1 SUMO protease to remove the SUMO tag, where required. Protein was then further purified by size exclusion with a Superdex™ 200 Increase 10/300 GL column (Cytiva) in gel filtration buffer (25 mM HEPES-NaOH pH 7.4, 200 mM NaCl, 5% glycerol, 5 mM 2-mercaptoethanol). Peak fractions were collected and analyzed using SDS PAGE, concentrated, snap-frozen in liquid nitrogen, and stored at −80°C.

To ^32^P-label proteins for EMSA and protein-protein crosslinking experiments, variants of ΔCARDS MDA5 and LGP2 having an engineered protein kinase A (PKA) site at the C-terminus (…RRASV), introduced by the Quik-Change II XL Site-Directed Mutagenesis Kit (Agilent) and primers listed in **Table S1**, were generated. Proteins were expressed and purified as described above. The purified proteins were ^32^P-labeled by incubation with [γ-^32^P]ATP (6000Ci/mmol, Revvity) and cAMP-dependent Protein Kinase (NEB) for 1 hour at room temperature in gel filtration buffer.

For site-specific protein-protein crosslinking, the artificial photo reactive amino acid p-benzoyl-l-phenylalanine (pBPA) was introduced into the LGP2-PKA construct by incorporating the amber codon (TAG) at indicated residues using the Quik-Change II XL Site-Directed Mutagenesis Kit (Agilent) and primers listed in **Table S1**. BLR(DE3) *E.coli* cells (Novagen) were co-transformed with LGP2-PKA and and pEVOL-pBpF (45)(Addgene, Plasmid #31190) plasmids and grown in filtered LB supplemented with 50 mM KH_2_PO_4_, 50 mM Na_2_HPO_4_, 25 mM (NH_4_)_2_SO_4_, 2 mM MgSO_4_, 0.5% glycerol, and 0.05% glucose. Protein expression and pBPA incorporation were induced by addition of 0.25 mM IPTG, 0.02% arabinose, and 1 mM pBPA (Fisher) at 16°C for 16 hours in the dark. pBPA-containing LGP2 proteins were purified and ^32^P-labeled as described above, with all steps performed in the dark to prevent UV exposure.

It is well established that the CARDs of MDA5 are not required for dsRNA binding and association with LGP2 (12, 24, 31). For this reason, ΔCARDs MDA5 constructs were used for subsequent biophysical and biochemical experiments.

### Mammalian cell culture

HEK293T cells (ATCC) were grown in Dulbecco’s Modified Eagle Medium (GenClone) supplemented with 10% heat-inactivated Bovine Calf Serum (HI-FBS, Hyclone) at 37°C with 5% CO_2_.

### Dual-luciferase reporter assay for IFN-β induction

Human MDA5/pUNO1 and LGP2/pUNO1 plasmids were purchased from Invivogen and mutations were introduced by the Quik-Change II XL Site-Directed Mutagenesis Kit (Agilent). Mutagenesis primers used in this study are listed in **Table S1**. For IFN-β induction assays, 500,000 cells were seeded in each well of a 24-well plate (Corning). After 24 hours, each well was transfected with 1 ng MDA5/pUNO1 plasmid, various concentrations of LGP2/pUNO1 plasmid or ΔCARDs MDA5/pUNO1 (as indicated), 6 ng pRLTK renilla luciferase reporter plasmid (Promega), and 150 ng IFN-β/Firefly luciferase reporter plasmid (Promega) using Lipofectamine 2000 transfection reagent (ThermoFisher Scientific). Cells were incubated for 24 hours before transfection with 1 ug/well of low molecular weight polyinosine-polycytidylic acid (LMW poly(I:C)), Invivogen) or synthetic dsRNA using Lipofectamine 2000 transfection reagent (ThermoFisher Scientific). After 18 hours, growth medium was aspirated, HEK293T cells were lysed, and IFN-β induction was measured using the Dual-Luciferase Reporter Assay System (Promega) and a Synergy Neo2 Hybrid Multi-Mode Reader with Gen5 software (Biotek). The IFN-β induction level was quantified as the firefly luciferase activity normalized to the renilla luciferase activity and analyzed with GraphPad Prism. Comparisons between more than two conditions, where indicated, were performed by one-way ANOVA in GraphPad Prism.

### In vitro transcription and dsRNA preparation

The longer dsRNAs used in this study (dsRNA150 and dsRNA300) were prepared by *in vitro* transcription and dsDNA templates were generated by PCR amplification of the pcDNA3.1 plasmid, with primers containing the T7 promoter sequence on the 5’ termini. The DNA templates for transcribing dsRNA50 were synthesized by IDT (Integrated DNA Technologies) and contain 2’o-methyl modifications on the first two nucleotides of the 5’ termini. The DNA template for transcribing pIC(150) were synthesized by BioSynthesis. Sequences of PCR primers and the transcribed RNAs used in this study are listed in **Table S1** and **Table S2,** respectively. *In vitro* transcription reactions (0.25 ml) were performed using 1 uM in-house-prepared T7 RNA polymerase (harboring a P266L mutation (52)) and 0.25 uM dsDNA template in a buffer containing 40 mM Tris-HCl pH 8.0, 10 mM NaCl, 30 mM MgCl_2_, 2 mM spermidine, 20 mM DTT, 100 U of RNaseOUT™ Recombinant Ribonuclease Inhibitor (Invitrogen), and 5 mM of each NTP. Reactions were incubated at 37°C for 2 hours and stopped by the addition of an equal volume of 95% formamide/0.05 M EDTA. Reaction products were resolved on denaturing PAGE gels (6 M urea) run in 1X TBE buffer: 15% PAGE for dsRNA50, 10% PAGE for dsRNA150, 6% PAGE for pIC(150), and 5% PAGE for dsRNA300. The desired products were then excised and eluted from the gel. The eluted RNA was ethanol precipitated, resuspended in an RNA storage buffer (6 mM Na-MOPS pH 6, 1 mM EDTA), and stored at −80°C until use.

Where indicated, the antisense RNA strand was 5’-labeled using [γ-^32^P]ATP (6000Ci/mmol, Revvity) and T4 polynucleotide kinase (NEB) and then annealed to the complementary sense strand. To anneal the dsRNAs, RNAs were diluted in buffer containing 30 mM HEPES-NaOH pH 7.4 and 10 mM potassium acetate, heated for 1 minute at 95°C, and cooled down (1°C/minute) to 25°C in a thermocycler. The purity of annealed dsRNAs was assessed by running samples on a 6% PAGE gel, staining with GelRed (Biotium), and visualizing products using a Bio-Rad ChemiDoc^TM^ imager.

### Electrophoretic mobility shift assays (EMSAs)

Protein-dsRNA complexes were assembled at room temperature in a buffer containing 40 mM Tris-HCl pH 8, 100 mM NaCl, 5% glycerol, 10 mM MgCl_2_, 10 mM 2-mercaptoethanol, and 0.1 mg/ml rAlbumin (NEB), and incubated with 20 nM dsRNA for 10 minutes at room temperature. To assess heteromeric complex formation, 20 nM dsRNA was first incubated with either LGP2 (10 minutes) followed by ΔCARDs MDA5 (20 minutes), or with ΔCARDs MDA5 first (20 minutes) followed by LGP2 (10 minutes). Protein concentrations are provided in the corresponding figure legends. Reactions utilizing dsRNA150 were resolved in 4% PAGE gels, while dsRNA50 reactions were resolved in 6% PAGE gels. All gels were run in 0.5X TBE buffer for 45 minutes at 100 V at 4°C. The products of the reactions were visualized by autoradiography using a PhosphorImager (Cytiva).

Protein-dsRNA300 complexes were assembled at room temperature in a buffer containing 40 mM Tris-HCl pH 8, 100 mM NaCl, 5% glycerol, 10 mM MgCl_2_, 10 mM 2-mercaptoethanol, and 0.1 mg/ml rAlbumin (NEB), and incubated with 100 nM dsRNA300 for 10 minutes at room temperature. To assess heteromeric complex formation, 50 nM dsRNA was first incubated with 1.5 uM LGP2 (10 minutes) followed by ΔCARDs MDA5 (20 minutes). Protein concentrations are provided in the corresponding figure legend. Where indicated, 4.5 mM AMP-PNP was added to the reaction. Reactions were resolved in 4% PAGE gels run in 0.5X TBE buffer for 45 minutes at 100 V at 4°C. The gels were then stained with GelRed (Biotium) for 5 minutes at room temperature and products were visualized using a Bio-Rad ChemiDoc^TM^ imager

### Mass photometry measurements

All manual dilution mass photometry data were recorded using a TwoMP (Refeyn Ltd) using MassGlass Uncoated (MGUC) microscope coverslips (Refeyn Ltd). Experiments were done in Normal mode with a Regular Field of View. Movies were collected for 60 seconds. Data was acquired and analyzed with AcquireMP and DiscoverMP software (Refeyn Ltd), respectively. The measured contrasts were converted to mass using the protein calibrant, MassFerence™ P1 Calibrant (Refeyn Ltd). For manual dilution measurements, samples were diluted in buffer consisting of 40 mM Tris-HCl pH 8, 100 mM NaCl, 10 mM MgCl_2_, and 10 mM 2-mercaptoethanol, similar to EMSA conditions. For measurements of individual protein and dsRNA components, samples were diluted 20-fold directly in a silicon gasket (Refeyn Ltd) prefilled with dilution buffer prior to data acquisition, resulting in a final concentration of 15 nM. Mass photometry measurements of dsRNA substrates were performed on glass slides treated with 0.01% poly-l-lysine (PLL, Sigma) to facilitate RNA surface immobilization.

Due to the high affinity of LGP2-dsRNA interactions (12, 25), LGP2-containing complexes remained intact following manual dilution, allowing standard mass photometry measurements to be performed. The LGP2-dsRNA150 and ΔCTT LGP2-dsRNA150 complexes were assembled by incubating 300 nM LGP2 with 20 nM dsRNA150 for 10 minutes at room temperature in EMSA buffer containing 40 mM Tris-HCl pH 8, 100 mM NaCl, 5% glycerol, 10 mM MgCl_2_, and 10 mM 2-mercaptoethanol. The LGP2-dsRNA300 complex was assembled by incubating 600 nM LGP2 with 20 nM dsRNA150 for 10 minutes at room temperature in EMSA buffer. The complexes were then diluted 20-fold directly in a silicon gasket (Refeyn Ltd) prefilled with dilution buffer prior to data acquisition to obtain final concentrations of 15 nM LGP2 and 1 nM dsRNA150 and 30 nM LGP2 and 1 nM dsRNA300.

### MassFluidix HC detection of MDA5 filaments

Manual dilution of MDA5 filaments to the nanomolar concentrations required for standard mass photometry measurements (53) resulted in the detection of only a small population of filament species, likely due to filament disassembly upon dilution. To preserve filament integrity and increase filament detection, complexes were rapidly diluted using the microfluidic MassFluidix HC system (MFx, Refeyn Ltd), which reduced the time between dilution and measurement to approximately 37 ms.

Rapid dilution measurements were performed on a TwoMP with the MFx system using uncoated MassFluidix HC (5-channel) chips (Refeyn Ltd) for the ΔCARDs MDA5-dsRNA150, LGP2-ΔCARDs MDA5-dsRNA150, LGP2-SUMO ΔCARDs MDA5-dsRNA150, ΔCTT LGP2-ΔCARDs MDA5-dsRNA150, ΔCARDs MDA5-dsRNA300, and LGP2-ΔCARDs MDA5-dsRNA300 complexes. To assemble protein complexes, 10 uM LGP2 was incubated with 1 uM dsRNA150 or 0.5 uM dsRNA300 in a buffer containing 25 mM HEPES-NaOH pH 7.4, 100 mM NaCl, 10 mM MgCl2, and 2 mM DTT for 10 minutes at room temperature. 10 uM ΔCARDs MDA5 was then added to the complex and incubated for 20 minutes at room temperature prior to overnight dialysis at 4°C in the same buffer. Complexes were then incubated at room temperature for 30 minutes prior to loading onto the MFx system. Measurements were collected using the MassFluidix module in AcquireMP (Refeyn Ltd) in Normal mode with a Large Field of View. Upon reaching the observation window, the sample dilution rate was set to 1,000x and complexes were rapidly diluted with Dulbecco’s phosphate-buffered saline (DPBS, Gibco). Movies were collected for 60 seconds and data were analyzed in DiscoverMP (Refeyn Ltd). MFx data were converted to mass using the protein calibrant, MassFerence™ P1 Calibrant (Refeyn, Ltd), measured by manual dilution.

### Mass photometry analysis of MDA5 filament dissociation kinetics

Consistent with the greater stability of MDA5 filaments on longer dsRNAs (40), a small population of intact filaments remained detectable on dsRNA300 following manual dilution, whereas no intact filaments were detected on dsRNA150. Consequently, dsRNA300 was selected for filament dissociation kinetics experiments to ensure sufficient detection of intact filaments following manual dilution. ΔCARDs MDA5-dsRNA300 and LGP2-ΔCARDs MDA5-dsRNA300 complexes were assembled at high concentrations and allowed to reach equilibrium, as described above. To initiate dissociation, complexes were first diluted 10-fold in dilution buffer and then diluted an additional 20-fold directly in a silicon gasket (Refeyn Ltd) prefilled with dilution buffer, yielding final concentrations of 50 nM ΔCARDs MDA5, 50 nM LGP2, and 2.5 nM dsRNA300. Complexes were mixed immediately prior to data acquisition and five consecutive 60-second movies were acquired for each complex in Normal mode with a Large Field of View. Data was acquired and analyzed with AcquireMP and DiscoverMP software (Refeyn Ltd), respectively. The measured contrasts were converted to mass using the protein calibrant, MassFerence™ P1 Calibrant (Refeyn Ltd).

The event counts of large filament species (800-2000 kDa) and total filament species (239-2000 kDa) were quantified at each time point using DiscoverMP software (Refeyn Ltd). The fraction of large filament species was calculated by dividing the number of large filament events by the total number of detected filament events. These values were plotted as a function of time in GraphPad Prism and fit to a one-phase exponential decay model. Dissociation kinetic parameters, including the half-life (t_1/2_), were determined from the fitted curves. Apparent dissociation rate constants (k_off_^app^) were then calculated according to the following equation: k_off_^app^ =0.693/t_1/2._

### Protein-protein crosslinking

For disuccinimidyl glutarate (DSG) crosslinking, protein complexes containing 200 nM LGP2 and/or 200 nM ΔCARDs MDA5 (unless otherwise indicated) were assembled in a buffer containing 40 mM HEPES-NaOH pH 7.4, 100 mM NaCl, 5% glycerol, 10 mM MgCl_2_, 10 mM 2-mercaptoethanol, and 0.1 mg/ml rAlbumin (NEB). 5 ug/ml high molecular weight polyinosine-polycytidylic acid (HMW poly(I:C), Invivogen) or 5 ug/ml dsRNA substrates were added last to the reaction, as indicated, and reactions incubated for 10 minutes at room temperature. Reactions were treated with DSG solution prepared in DMSO (ThermoFisher Scientific), resulting in a final DSG concentration of 0.4 mM, for 30 minutes at room temperature and stopped by the addition of an equal volume of solution containing 100 mM Tris-HCl pH 8, 2X Laemmli Sample Buffer (BioRad), and 10% 2-mercaptoethanol. Crosslinked products were resolved using 6% Tris-glycine SDS PAGE and visualized by autoradiography using a PhosphorImager (Cytiva).

The purified pBPA-containing LGP2 proteins were ^32^P-labeled as described above. Protein complexes containing 100 nM ^32^P-labeled LGP2-pBPA and 100 nM ΔCARDs MDA5 were assembled in a buffer containing 40 mM HEPES-NaOH pH 7.4, 100 mM NaCl, 5% glycerol, 10 mM MgCl_2_, 10 mM 2-mercaptoethanol, and 0.1 mg/ml rAlbumin (NEB). 1 ug/ml HMW pIC (Invivogen) was added to the reaction last and incubated for 10 minutes at room temperature. Crosslinking was activated by UV irradiation at 360 mn for 20 minutes at room temperature and stopped by the addition of an equal volume of solution containing 100 mM Tris-HCl pH 8, 2X Laemmli Sample Buffer (BioRad), and 10% 2-mercaptoethanol. Crosslinked products were resolved using 6% Tris-glycine SDS PAGE and visualized by autoradiography using a PhosphorImager (Cytiva).

### ATP hydrolysis assay

ATP hydrolysis assays were carried out using LGP2 (300 nM) in a buffer containing 40 mM Tris-HCl pH 8, 50 mM NaCl, 5% glycerol, 10 mM MgCl_2_, 10 mM 2-mercaptoethanol, 0.1 mg/ml rAlbumin (NEB), and 0.125 uM [γ-^32^P]ATP (6000Ci/mmol, Revvity). ATP hydrolysis was stimulated by the addition of 5 ug/ml HMW poly(I:C) (Invivogen) and reactions were carried out for the indicated time periods at room temperature and stopped by the addition of an equal volume of 95% formamide/0.05 M EDTA. The reactions were resolved by 20% PAGE containing 6 M urea and visualized by autoradiography using a PhosphorImager (Cytiva).

### Reproducibility

All *in vitro* experiments were performed at least three times independently with similar results. Representative data are shown. For cellular experiments, data are derived from at least three independent biological replicates. The number of independent replicates and statistical analyses used for each experiment are indicated in the corresponding figure legends.

## Author Contributions

G.B. and A.M.P. designed research; G.B., L.X., and Y-C.S. performed research; G.B. analyzed data; G.B. and A.M.P. wrote the paper.

## Acknowledgments

A.M.P. is an Investigator, G.B. is a Postdoctoral Scientist, and L.X. was a Research Specialist of the Howard Hughes Medical Institute (HHMI). We thank Pyle lab members for thoughtful comments and productive discussion. We thank the following mass photometry facilities: Yale West Campus Analytical Core and MIT Biophysical Instrumentation Facility. Specifically, we thank Dr. Terence Wu, Dr. Jandi Kim, and Dr. Ky Lowenhaupt. We also thank Kayleigh Fay and Dr. Matt Ranaghan from Refeyn for their guidance, expertise, and technical support in the design and analysis of mass photometry experiments.

