## Supporting Information for "Heteromeric LGP2-MDA5 receptor complexes drive antiviral signaling"

### Supporting Information for
Heteromeric LGP2-MDA5 receptor complexes drive antiviral signaling

Gina Buchel[1,2], Ling Xu[1,2,†], Yu-Chieh Sung[1], and Anna Marie Pyle[1,2,3]*

[1]Department of Molecular, Cellular, and Developmental Biology, Yale University, New Haven, CT 06511

[2]Howard Hughes Medical Institute, Chevy Chase, MD 20815

[3]Department of Chemistry, Yale University, New Haven, CT 06511

[†]Current address: Department of Chemistry and RNA Center, The Herbert Wertheim University of Florida Scripps Institute for Biomedical Innovation and Technology, Jupiter, FL 33458

**This PDF file includes:**

Figures S1 to S12
Tables S1 to S3
SI References

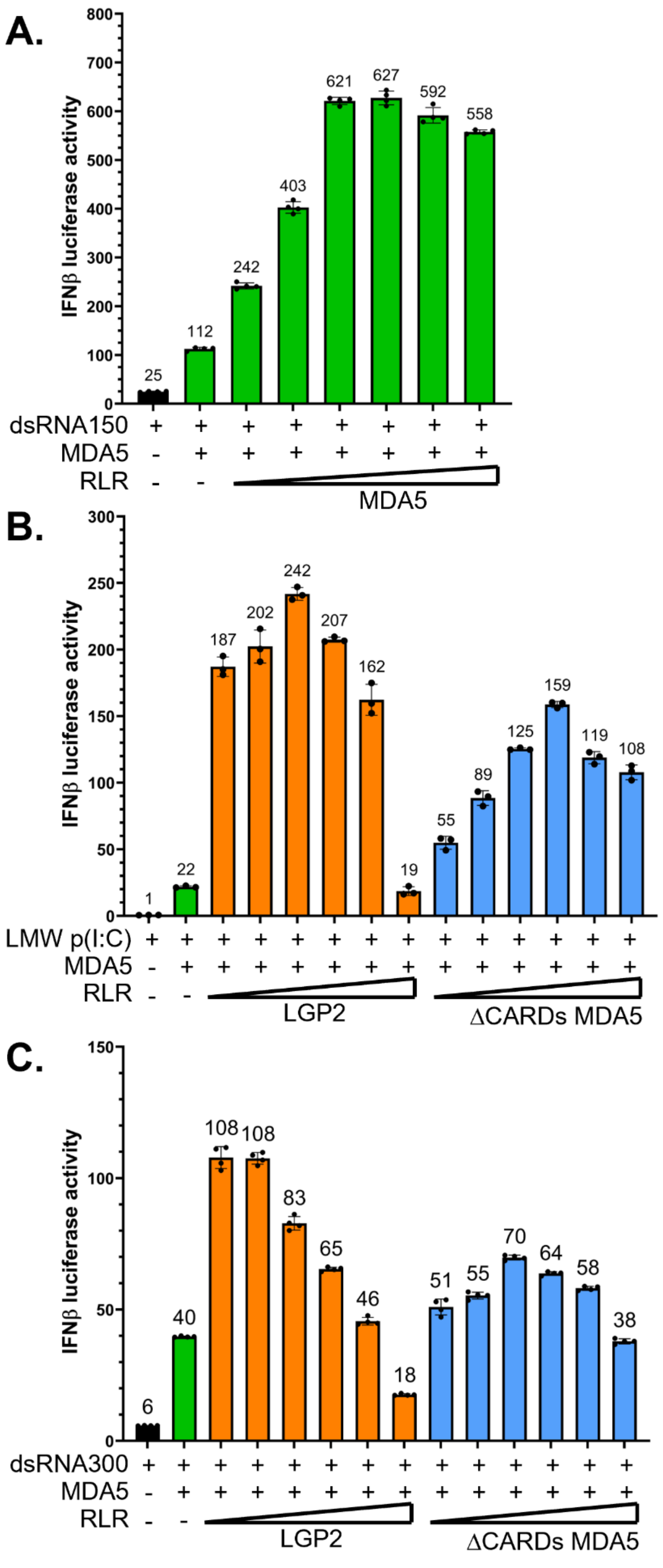
A.
IFNβ luciferase activity
0
100
200
300
400
500
600
700
800
25
112
242
403
621
627
592
558
dsRNA150
MDA5
RLR
MDA5
B.
IFNβ luciferase activity
0
50
100
150
200
250
300
1
22
187
202
242
207
162
19
55
89
125
159
119
108
LMW p(I:C)
MDA5
RLR
LGP2
ΔCARDs MDA5
C.
IFNβ luciferase activity
0
50
100
150
6
40
108
108
83
65
46
18
51
55
70
64
58
38
dsRNA300
MDA5
RLR
LGP2
ΔCARDs MDA5

**Fig. S1. RLR-mediated induction of IFN signaling by dsRNAs.**
**A**. IFN induction by co-transfection of MDA5 (1 ng) and additional MDA5 (5, 10, 20, 50, 100, 200 ng) plasmid stimulated with 1 ug dsRNA150 in HEK293T cells. Data are represented as mean±SD (n=4 biological replicates).
**B**. IFN induction by co-transfection of MDA5 (1 ng) and LGP2 (orange) or ∆CARDs MDA5 (blue) plasmids (5, 10, 20, 50, 100, 200 ng) stimulated with 1 ug LMW poly(I:C) in HEK293T cells. Data are represented as mean±SD (n=3 biological replicates).
**C**. IFN induction by co-transfection of MDA5 (1 ng) and LGP2 (orange) or ∆CARDs MDA5 (blue) plasmids (5, 10, 20, 50, 100, 200 ng) stimulated with 1 ug dsRNA300 in HEK293T cells. Data are represented as mean±SD (n=4 biological replicates).

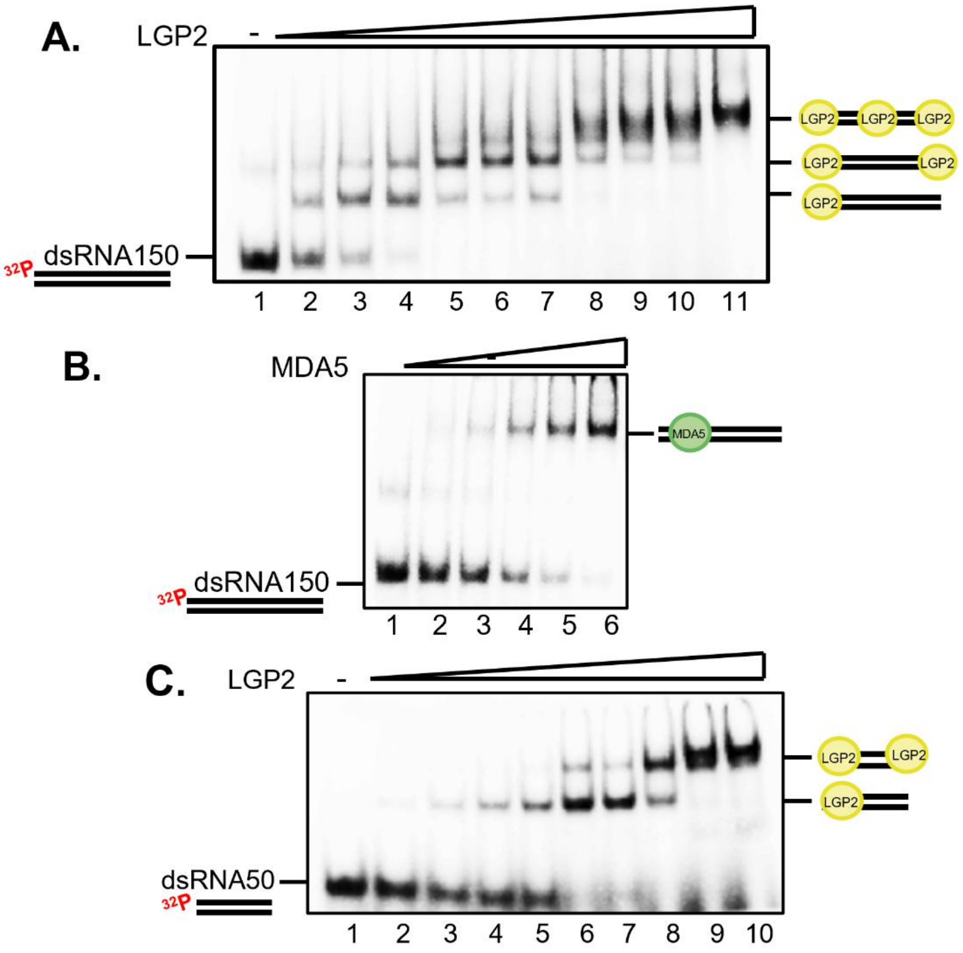


**Fig. S2. Binding of individual components to dsRNA150 and dsRNA50.**

**A, B.** EMSA performed using 20 nM dsRNA150-$^{32}$P with increasing amounts of (**A**) LGP2 (40, 80, 120, 160, 200, 220, 240, 260, 280, 300 nM) or (**B**) MDA5 (100, 150, 200, 250, 300 nM).

**C**. EMSA performed using 20 nM dsRNA50-$^{32}$P with increasing amounts of LGP2 (40, 80, 100, 120, 160, 180, 200, 240, 300 nM).

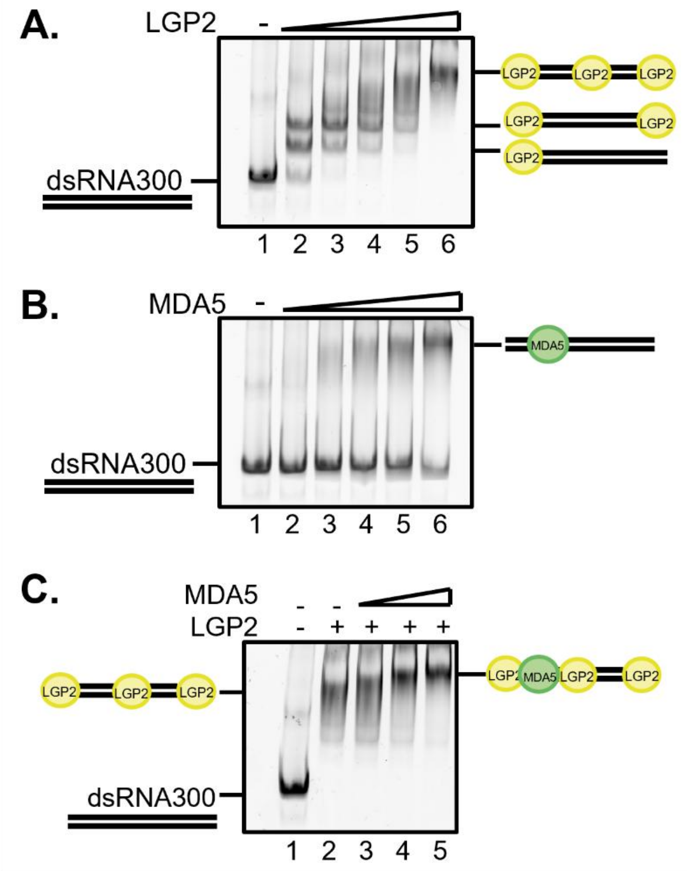


**Fig. S3. Binding of individual components to dsRNA300 and heteromeric complex assembly.**
**A, B.** EMSA performed using 100 nM dsRNA300 with addition of increasing concentrations (0.5, 1.0, 1.5, 2,0, 3.0, uM) of either (**A**) LGP2 or (**B**) MDA5. Gels were stained with GelRed.
**C**. EMSA performed using 50 nM dsRNA300 with a pre-existing LGP2-dsRNA300 complex (where 1.5 uM LGP2 was pre-loaded) and increasing amounts of MDA5 were added to the complex (0.5, 1.0, 1.5 uM).

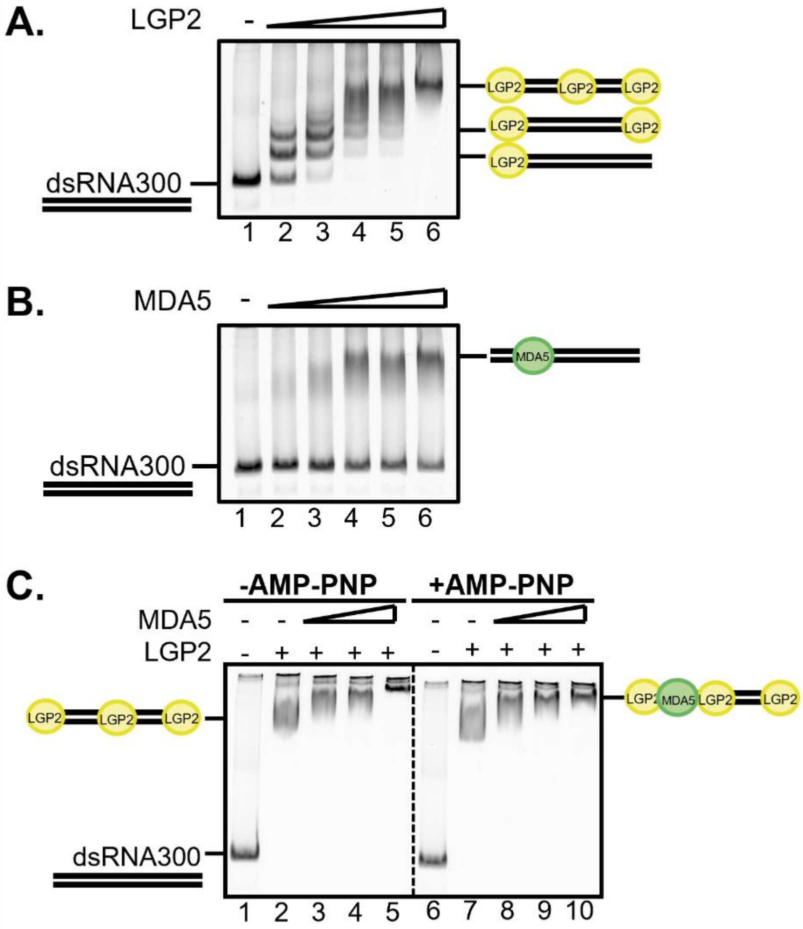


**Fig. S4. Binding of individual components to dsRNA300 and heteromeric complex assembly in the presence of AMP-PNP.**

**A, B.** EMSA performed using 100 nM dsRNA300 with addition of increasing concentrations (0.5, 1.0, 1.5, 2,0, 3.0, uM) of either (**A**) LGP2 or (**B**) MDA5 in the presence of 4.5 mM AMP-PNP. Gels were stained with GelRed.

**C**. EMSA performed using 50 nM dsRNA300 with a pre-existing LGP2-dsRNA300 complex (where 1.5 uM LGP2 was pre-loaded) and increasing amounts of MDA5 were added to the complex (0.5, 1.0, 1.5 uM) in the absence (Lanes 1-5) and presence of 4.5 mM AMP-PNP (Lanes 6-10).

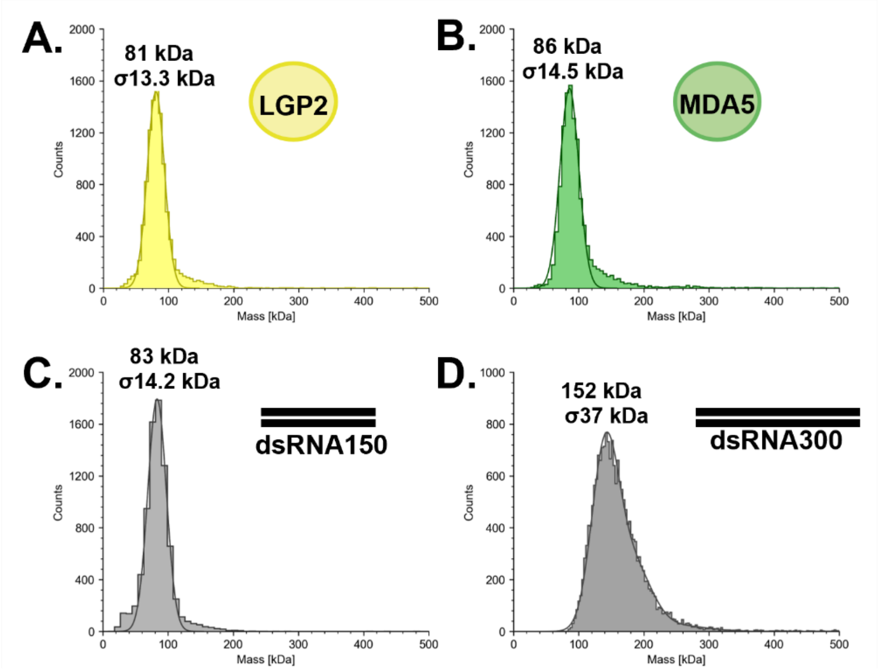


**Fig. S5. Mass photometry measurements for heteromeric complex components.**
**A-D.** Mass photometry measurements for individual components following manual 20-fold droplet dilution resulting in the final concentrations: (**A**) 15 nM LGP2 (~81 kDa, yellow), (**B**) 15 nM MDA5 (~87 kDa, green), (**C**) 15 nM dsRNA150 (~83 kDa, light grey), and (**D**) 15 nM dsRNA300 (~152 kDa, dark grey). Slides were coated with 0.01% PLL for measurements of dsRNA150 and dsRNA300 (**C**,**D**). Gaussian fitting is shown.

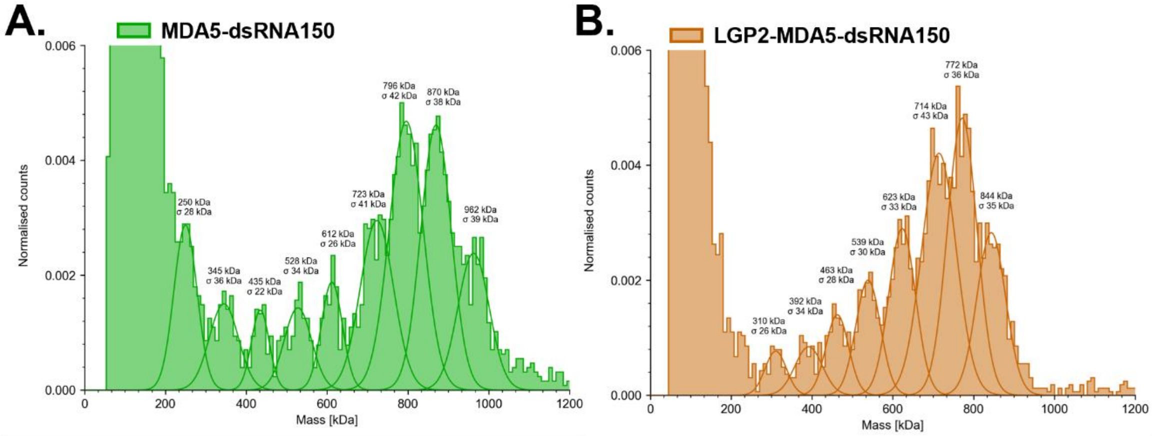


**Fig. S6. Mass photometry measurements for heteromeric complex assembly on dsRNA150.** **A**, **B.** Mass photometry measurements for (**A**) MDA5-dsRNA150 and (**B**) LGP2-MDA5-dsRNA150 complexes utilizing the MassFluidix HC microfluidics system (Refeyn Ltd). Gaussian fitting is shown with quantification of areas under the curve to evaluate filament stoichiometric abundance.

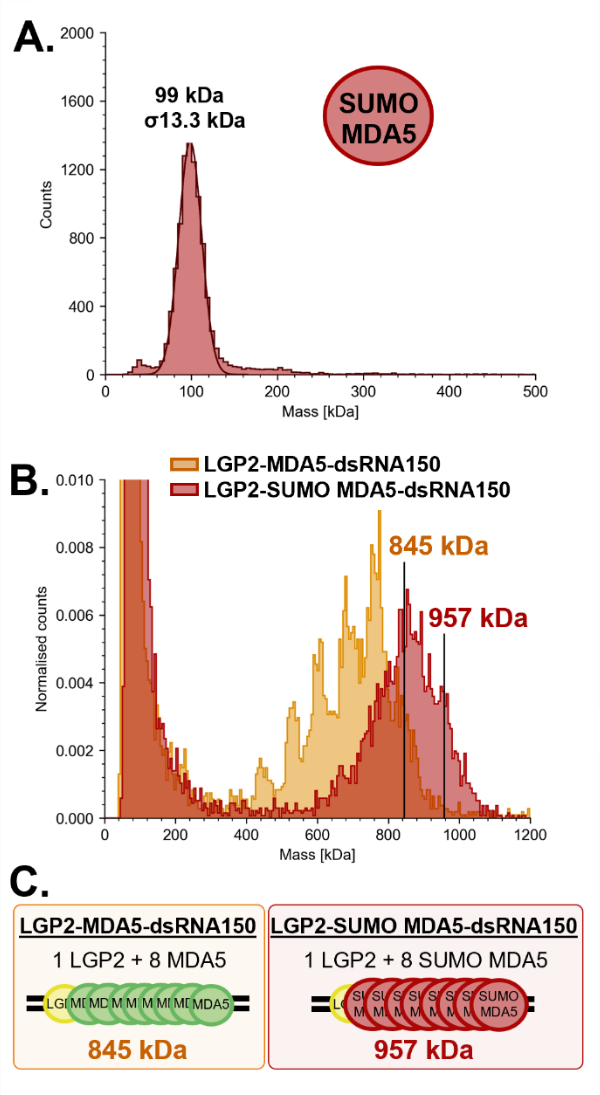


**Fig. S7. Mass photometry measurements for SUMO-tagged MDA5 and heteromeric complex assembly with LGP2.**

**A**. Mass photometry measurement following manual 20-fold droplet dilution resulting in the final concentration of 15 nM SUMO-MDA5 (~99 kDa, red). Gaussian fitting is shown.

**B**. Mass photometry measurements for LGP2-MDA5-dsRNA150 (orange), and LGP2-SUMO-MDA5-dsRNA150 (red) complexes utilizing the MassFluidix HC microfluidics system (Refeyn Ltd). Molecular weight markers represent the median molecular weight of the largest filament length measured for comparison.

**C**. Determination of filament stoichiometry based on addition of a SUMO tag (~13 kDa) to MDA5.

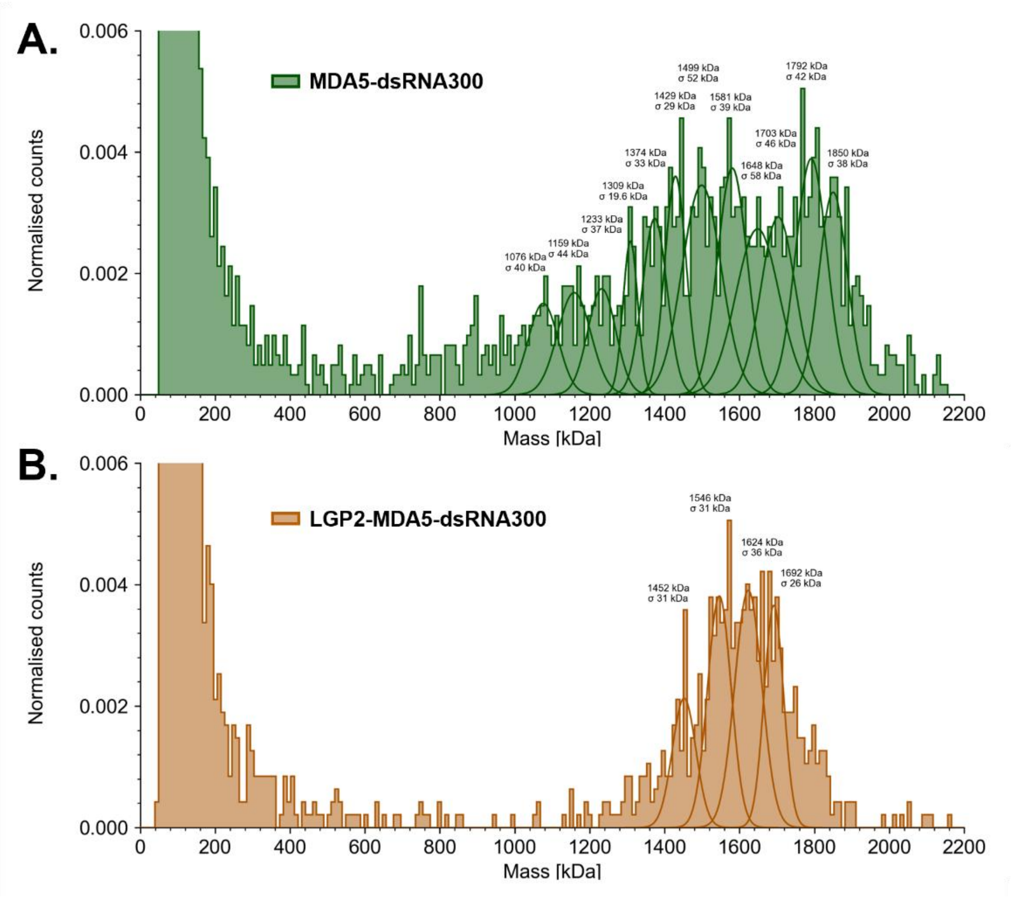


**Fig. S8. Mass photometry measurements for heteromeric complex assembly on dsRNA300.** **A**, **B**. Mass photometry measurements for (**A**) MDA5-dsRNA300 and (**B**) LGP2-MDA5-dsRNA300 complexes utilizing the MassFluidix HC microfluidics system (Refeyn Ltd). Gaussian fitting is shown with quantification of areas under the curve to evaluate filament stoichiometric abundance.

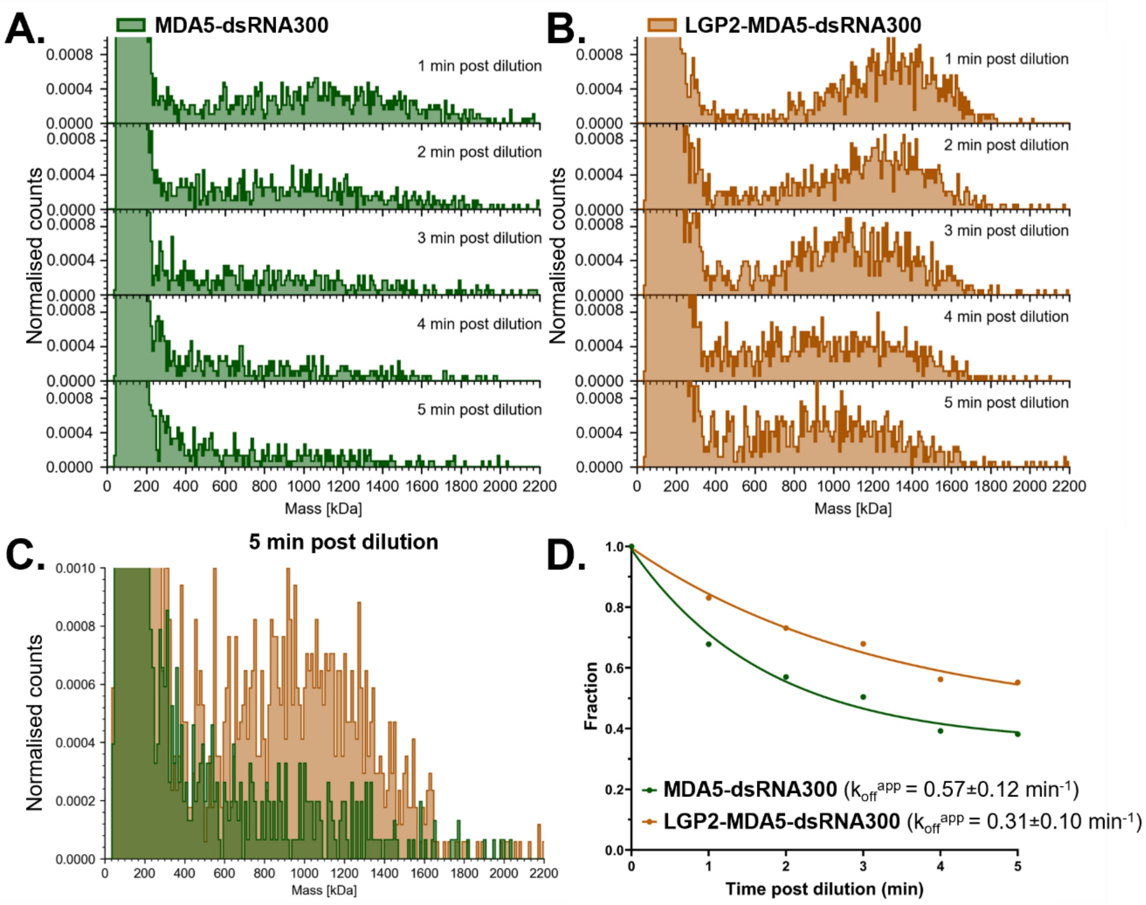


**Fig. S9. LGP2 increases MDA5 filament stability.**

**A, B.** Mass photometry measurements for (**A**) MDA5-dsRNA300 (dark green) and (**B**) LGP2-MDA5-dsRNA300 (dark orange) complexes following manual dilution (1:200 dilution) over time. Five consecutive one-minute measurements were acquired following manual dilution directly within the silicone gasket on the coverslip. Complexes were mixed between measurements to ensure consistent counts throughout the experiment. Unbound proteins (~79-85 kDa) are excluded from the analysis.

**C.** Overlay of mass photometry histograms for MDA5-dsRNA300 (dark green) and LGP2-MDA5-dsRNA300 (dark orange) complexes at the five-minute post dilution time point to show relative abundance of filaments.

**D.** Relative abundance of large filaments ($\geq$800 kDa) following manual dilution. The fraction of large filaments was monitored over time and fit to a one-phase exponential decay model in GraphPad Prism for MDA5-dsRNA300 (dark green, $R^2$= 0.99) and LGP2-MDA5-dsRNA300 (dark orange, $R^2$= 0.99). Dissociation kinetics were determined from the fitted curves, including the half-life ($t_{1/2}$) and standard error values. Apparent dissociation rate constants ($k_{off}^{app}$) were calculated using $k_{off}^{app}$ =0.693/$t_{1/2}$ and are displayed on the plot.

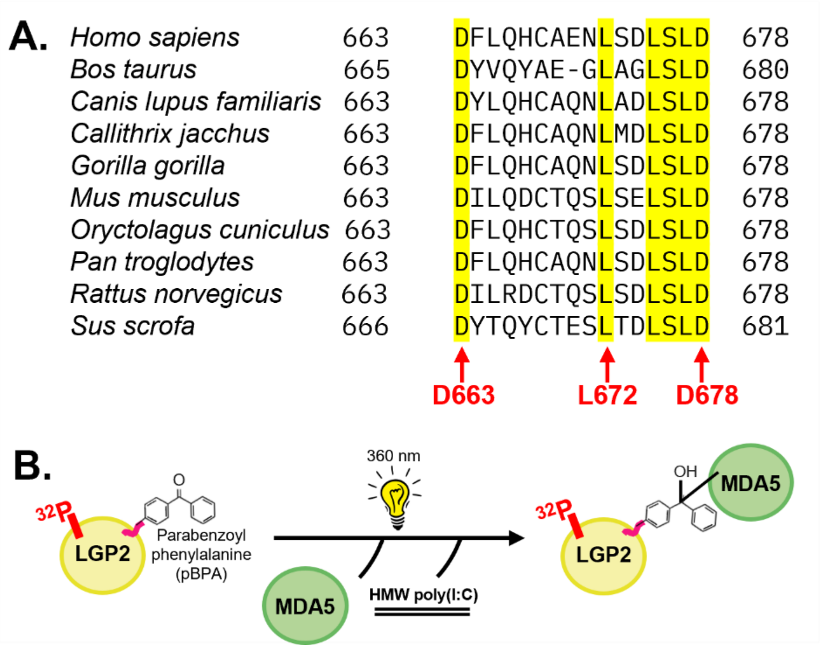


**Fig. S10. Conservation analysis of the CTT of LGP2 and pBPA incorporation.**
**A**. Conservation of the CTT of LGP2 (residues 663-678 in *Homo sapiens*) across mammalian species. Highlighted residues indicate 100% homology across all listed species. The red arrow indicates the residues where pBPA was incorporated (D663, L672, D678) by an *E. coli* expression system (1). Sequences were obtained from NCBI (**Table S3**) and aligned using Clustal Omega (2).
**B**. Schematic representation showing incorporation of a site-specific photo-crosslinking probe, parabenzoyl phenylalanine (pBPA), into the CTT (residues D663, L672, or D678) of $^{32}$P-labeled LGP2 for protein-protein crosslinking experiments. Experiments were performed in the presence of MDA5 and HMW poly(I:C) and subjected to UV (360 nm) for 20 minutes at room temperature.

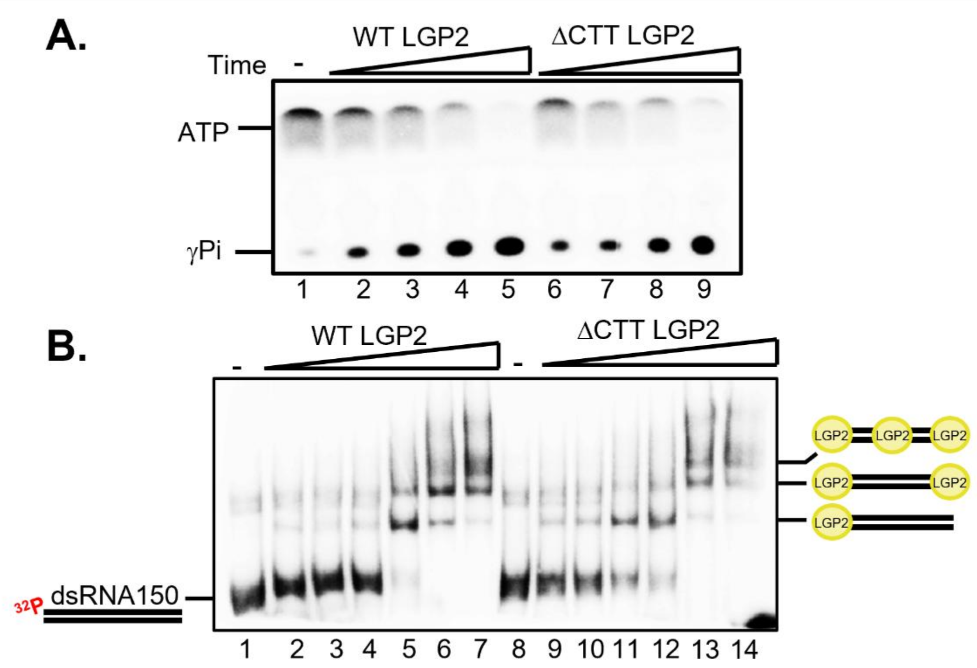


**Fig. S11. Deletion of the CTT of LGP2 (residues 663-678) does not impair ATPase or dsRNA-binding activities of the enzyme.**

**A**. ATP hydrolysis assay was performed using [γ-$^{32}$P]ATP in the presence of 300 nM WT LGP2 (Lanes 2-5) or 300 nM ΔCTT LGP2 (Lanes 6-9). Experiments were performed in the presence of 5 ug/ml HMW poly(I:C) to stimulate LGP2 ATPase activity. Reactions were carried out at RT for 5, 10, 20, 40 minutes.

**B**. EMSA was performed using 20 nM dsRNA150-$^{32}$P with increasing amounts of WT (Lanes 2-7) or ΔCTT (Lanes 9-14) LGP2 (80, 120, 160, 200, 260, 300 nM).

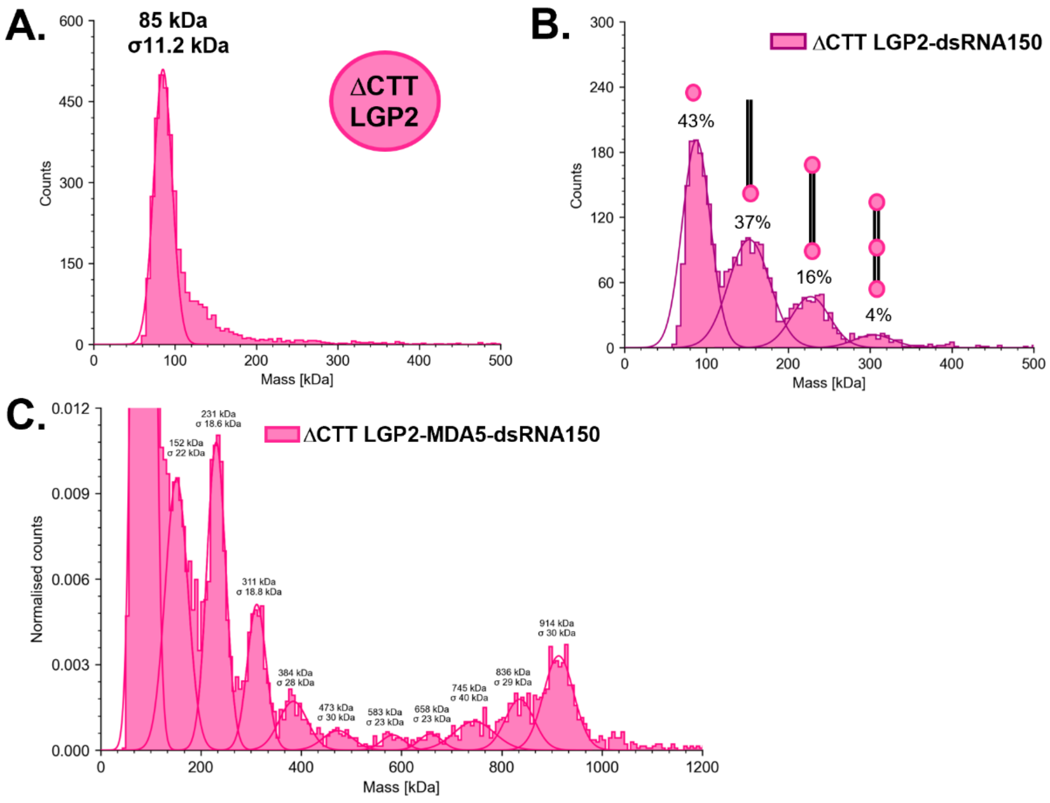


**Fig. S12. Mass photometry measurements for ΔCTT LGP2 and dsRNA150-containing complexes.**

**A**. Mass photometry measurement following manual 20-fold droplet dilution, yielding a final concentration of 15 nM ΔCTT LGP2 (~85 kDa). Gaussian fitting is shown.

**B**. Mass photometry measurement for the LGP2-dsRNA150 complex following manual 20-fold droplet dilution. Molecular weights correspond to free ΔCTT LGP2 (~88 kDa), ΔCTT LGP2-dsRNA150 (~153 kDa), 2ΔCTT LGP2-dsRNA150 (~227 kDa), and 3ΔCTT LGP2-dsRNA150 (~304 kDa). Gaussian fitting is shown.

**C**. Mass photometry measurements for the ΔCTT LGP2-MDA5-dsRNA150 complex utilizing the MassFluidix HC microfluidics system (Refeyn Ltd). Gaussian fitting is shown with quantification of areas under the curve to evaluate filament stoichiometric abundance.

**Table S1. Primers used in this study.**

| Primer name | Sequence (5' – 3') |
|---|---|
| **Mutagenesis primers** | |
| dCARDs MDA5 pUNO_FWD | GCAAGAGCATCCCCGGAGCCAGAAC |
| dCARDs MDA5 pUNO_REV | CATGTTGACCGGTGATCTCAGGTAG |
| MDA5 PKA pET28_FWD | GTTTATTTAGTGATGAGGATCGCAGGGCAAGTGTTTAGAGACA AGCTTAGGTATT |
| MDA5 PKA pET28_REV | AATACCTAAGCTTGTCTCTAAACACTTGCCCTGCGATCCTCAT CACTAAATAAAC |
| LGP2 PKA pET28_FWD | TGTCGGACCTCTCCCTGGACCGCAGGGCAAGTGTTTGAAGAC AAGCTTAGGTATT |
| LGP2 PKA pET28_REV | AATACCTAAGCTTGTCTTCAAACACTTGCCCTGCGGTCCAGGG AGAGGTCCGACA |
| LGP2 D663BPA pET28_FWD | TCCGTGCCTGACTTTTAGTTCCTGCAGCATTGT |
| LGP2 D663BPA pET28_REV | ACAATGCTGCAGGAACTAAAAGTCAGGCACGGA |
| LGP2 L672BPA pET28_FWD | CATTGTGCCGAGAACTAGTCGGACCTCTCCCTG |
| LGP2 L672BPA pET28_REV | CAGGGAGAGGTCCGACTAGTTCTCGGCACAATG |
| LGP2 D678BPA pET28_FWD | TCGGACCTCTCCCTGTAGCGCAGGGCAAGTGTT |
| LGP2 D678BPA pET28_REV | AACACTTGCCCTGCGCTACAGGGAGAGGTCCGA |
| LGP2 d663-678 pET28_FWD | CCTTCTCCGTGCCTGACTTTTGAAGACAAGCTTAGGTATT |
| LGP2 d663-678 pET28_REV | AATACCTAAGCTTGTCTTCAAAAGTCAGGCACGGAGAAGG |
| LGP2 d663-678 pUNO_FWD | CCTTCTCCGTGCCTGACTTTTGACCACCTCATTGCTGCTA |
| LGP2 d663-678 pUNO_REV | TAGCAGCAATGAGGTGGTCAAAAGTCAGGCACGGAGAAGG |
| **Primers for amplification of dsDNA templates by PCR** | |
| pcDNA3.1sense_T7FWD | TAATACGACTCACTATAGGGAGACCC |
| pcDNA3.1antisense_REV | GGGAGACCCAAGCTTGAATTCT |
| pcDNA3.1sense_150REV | GGAACAGATGGCTGGCAACTAGAA |
| pcDNA3.1antisense150_T7FWD | TAATACGACTCACTATAGGAACAGATGGCTGGCAACTAGAA |
| pcDNA3.1sense_300REV | GGATCCTCCCCCTTGCTGTCC |
| pcDNA3.1antisense_300T7FWD | TAATACGACTCACTATAGGATCCTCCCCCTTGCTGTCC |
| **DNA templates for dsRNA50 (containing 2' o-methyl at 5' ends)** | |
| RNA50 sense template | mGmGCTACACGAAAGCTCCAATTGTATGCCAGGGTACATCAT GCTGCGCACCTATAGTGAGTCGTATTA |
| RNA antisense template | mGmGTGCGCAGCATGATGTACCCTGGCATACAATTGGAGCTT TCGTGTAGCCTATAGTGAGTCGTATTA |
| **DNA templates for pIC(150)** | |
| RNA pIC(150) template | GGAGACCCCCGGGGGCCCCCGGGGGCCCCCGGGGGCCCCC GGGGGCCCCCGGGGGCCCCCGGGGGCCCCCGGGGGCCCCC CGGGGGCCCCCGGGGGCCCCCGGGGGCCCCCGGGGGCCC CCGGGGGCCCCCGGGGGCCCCCGGGGGTCTCCTATAGTGAG TCGTATTA |
| **T7 promoter for ssDNA templates** | |
| T7 promoter | TAATACGACTCACTATA |

**Table S2. Sequences of RNA used in this study.**

| RNA name | Sequence (5' – 3') |
|---|---|
| RNA150 - Sense | GGGAGACCCAAGCUUGAAUUCUGCAGAUAUCCAUCACACUGGCGGCC GCUCGAGCAUGCAUCUAGAGGGCCCUAUUCUAUAGUGUCACCUAAAU GCUACCGCUCGCUGAUCAGCCUCGACUGUGCCUUCUAGUUGCCAGC CAUCUGUUCC |
| RNA150 - Antisense | GGAACAGAUGGCUGGCAACUAGAAGGCACAGUCGAGGCUGAUCAGC GAGCGGUAGCAUUUAGGUGACACUAUAGAAUAGGGCCCUCUAGAUG CAUGCUCGAGCGGCCGCCAGUGUGAUGGAUAUCUGCAGAAUUCAAG CUUGGGUCUCCC |
| RNA50 - Sense | GGUGCGCAGCAUGAUGUACCCUGGCAUACAAUUGGAGCUUUCGUGU AGCC |
| RNA50 - Antisense | GGCUACACGAAAGCUCCAAUUGUAUGCCAGGGUACAUCAUGCUGCGC ACC |
| RNA300 - Sense | GGGAGACCCAAGCUUGAAUUCUGCAGAUAUCCAUCACACUGGCGGCC GCUCGAGCAUGCAUCUAGAGGGCCCUAUUCUAUAGUGUCACCUAAAU GCUACCGCUCGCUGAUCAGCCUCGACUGUGCCUUCUAGUUGCCAGC CAUCUGUUGUUUGCCCCUCCCCCGUGCCUUCCUUGACCCUGGAAGG UGCCACUCCCACUGUCCUUUCCUAAUAAAAUGAGGAAAUUGCAUCGC AUUGUCUGAGUAGGUGUCAUUCUAUUCUGGGGGGUGGGGUGGGGC AGGACAGCAAGGGGGAGGAUCC |
| RNA300 - Antisense | GGAUCCUCCCCCUUGCUGUCCUGCCCCACCCCACCCCCCAGAAUAGA AUGACACCUACUCAGACAAUGCGAUGCAAUUUCCUCAUUUUAUUAGG AAAGGACAGUGGGAGUGGCACCUUCCAGGGUCAAGGAAGGCACGGG GGAGGGGCAAACAACAGAUGGCUGGCAACUAGAAGGCACAGUCGAG GCUGAUCAGCGAGCGGUAGCAUUUAGGUGACACUAUAGAAUAGGGC CCUCUAGAUGCAUGCUCGAGCGGCCGCCAGUGUGAUGGAUAUCUGC AGAAUUCAAGCUUGGGUCUCCC |
| RNA pIC(150) | GIAIAIIIIICCCCCIIIIICCCCCIIIIICCCCCIIIIICCCCCIIIIICCCCCIIII ICCCCCIIIIICCCCCIIIIICCCCCIIIIICCCCCIIIIICCCCCIIIIICCCCCIIIIICCCCC IIIIICCCCCUCUCC |

**Table S3. LGP2 protein sequences used for sequence alignment.**

| Organism | NCBI accession number |
|---|---|
| *Homo sapiens* | NP_077024.2 |
| *Bos taurus* | NP_001015545.1 |
| *Canis lupus familiaris* | XP_038532413.1 |
| *Callithrix jacchus* | XP_054112007.1 |
| *Gorilla gorilla* | XP_055244089.1 |
| *Mus musculus* | NP_084426.2 |
| *Oryctolagus cuniculus* | XP_002719437.3 |
| *Pan troglodytes* | XP_016787279.1 |
| *Rattus norvegicus* | NP_001092258.1 |
| *Sus scrofa* | NP_001186061.1 |
